# Large-scale whole-genome sequencing of migratory Bogong moths *Agrotis infusa* reveals genetic variants associated with migratory direction in a panmictic population

**DOI:** 10.1101/2022.05.27.493801

**Authors:** Jesse RA Wallace, Ryszard Maleszka, Eric J Warrant

## Abstract

One of the most interesting macroscopic phenomena in the animal world is seasonal migration. A central goal of research into animal migration is to better understand the mechanisms that evolved to solve the complex challenges which a migratory life history presents. Each year, and with a high degree of species-level site fidelity, the Australian Bogong moth makes a return migration of up to and over 1000 km between widely distributed breeding grounds and a specific set of aestivation sites in the Australian Alps. It does this without any opportunity to learn the migratory route or the location of the aestivation sites from either older generations or repeated migrations, meaning that the information required by the moth to navigate during its migration must be inherited. The migratory direction, and therefore the inherited navigational information in Bogong moths, varies with breeding site, providing us with an opportunity to search for the source of that heritability by comparing the genomes of moths collected from different breeding areas. We successfully sequenced whole nuclear genomes of 77 Bogong moths collected from across their breeding grounds and summer range, and found that the Bogong moth population contains a large amount of (mostly rare) variation. We found no evidence of population structure, indicating that Bogong moths are panmictic. A genome-wide scan for signals of selection indicate that the Bogong population has recently recovered from a past bottleneck, however genomic regions which have likely undergone balancing selection were also detected. Despite panmixia, four genetic variants in breeding-ground-caught Bogong moths were found to be significantly associated with geographic location, and therefore migratory direction, indicating promising future avenues of research into the molecular basis of long-distance navigation.

## 1 Introduction

Migratory behaviour is one of the most conspicuous and fascinating phenomena in the natural world (Dingle, 2014). It has evolved independently in many taxa across the animal kingdom, despite frequently requiring complex physiological and neurological adaptations for successful execution (Chapman et al., 2015). These adaptations are particularly pronounced in animals that navigate over great distances from a distinct origin to a specific destination, and that return to the origin after the season changes, especially if the origin and/or destination remain stable for an individual or population from one year to the next. For such fidelity to be possible, the migrant must either learn its migratory destination or inherit the information required to find it.

The Australian Bogong moth *Agrotis infusa* is a wonderful example of an animal that migrates in an extraordinarily directed and precise manner (Fig. 1) (reviewed by Warrant et al., 2016). The migratory journey of the adult Bogong moth starts in their breeding grounds, the dry plains of southern Queensland, western NSW, western Victoria, and eastern South Australia (Common, 1954; Warrant et al., 2016), a vast arc of country that spans at least 7° of latitude and 9° of longitude—1,400 km from end to end. In spring, the moth leaves its breeding area and flies up to 1,000 km to a specific set of sites located in the mountaintops of the Australian Alps, a narrow strip of alpine territory stretching between Mt. Gingera in the north and Mt. Buller in the south, a summer range which spans less than 2° of latitude and 2° of longitude, or some 274 km from end to end. Thus, the Bogong moth’s breeding grounds are vast relative to their summer range. This means that the required direction of migration must vary, depending on the starting point in the breeding grounds (*arrows* in Fig. 1) (Warrant et al., 2016). For instance, moths migrating to the Australian Alps from southern Queensland must fly south, whereas moths migrating to the Australian Alps from eastern South Australia must fly east.

**Figure 1:**
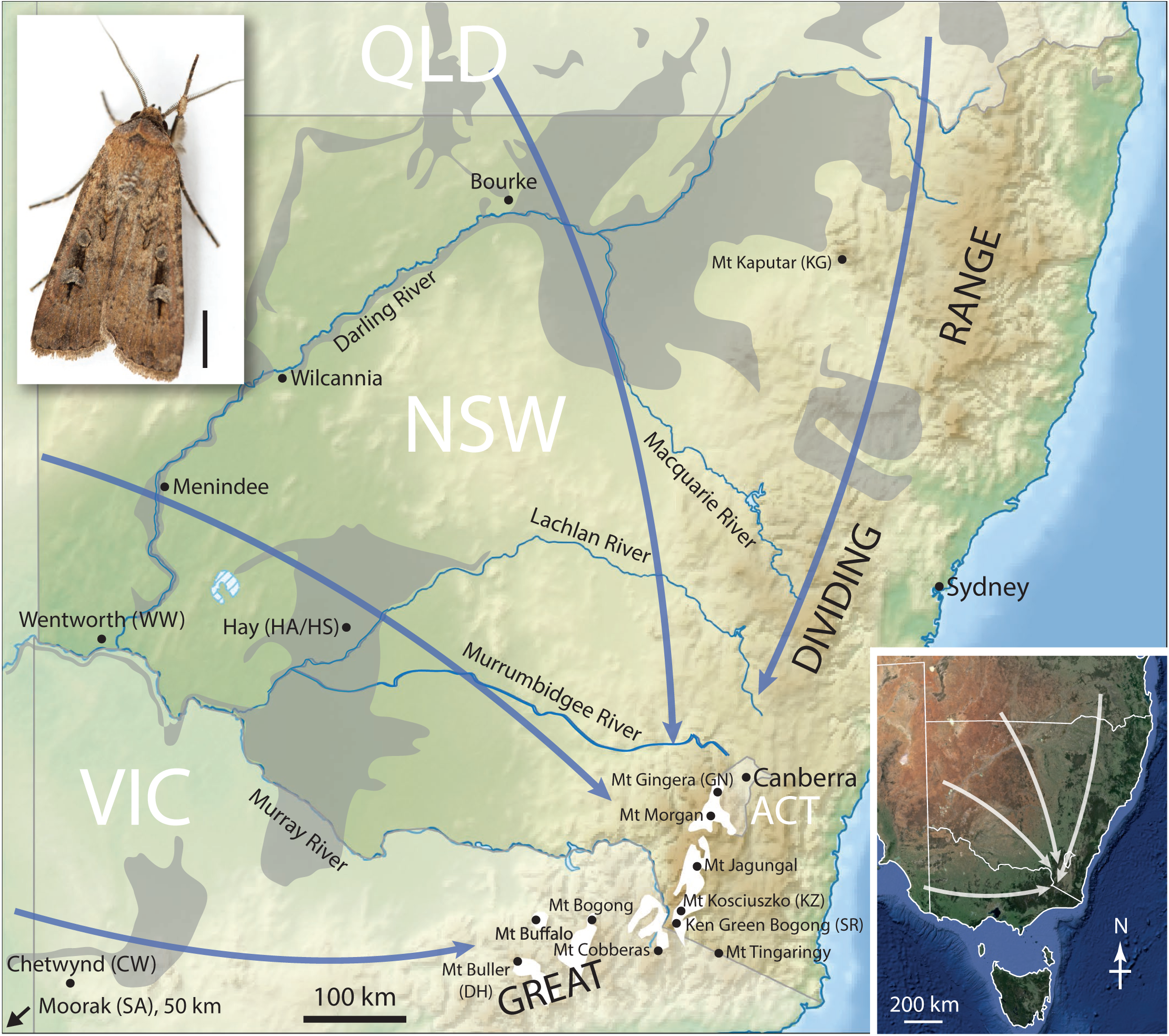
Map of Bogong moth migratory routes showing the names of the major rivers in New South Wales (*NSW*), and the locations of various towns and mountain peaks, mentioned in the text (shown in the context of continental south-eastern Australia: *inset, lower right*). Areas of grey cracking clays—favoured soils for Bogong moth winter development—are shown in *grey*. Bogong moths fly in different directions (*arrows*) towards the Australian Alps from various regions of south-eastern Australia, from as far distant as eastern South Australia, western Victoria (*VIC*), western and north-western New South Wales, and southern and south-eastern Queensland (*QLD*). The *white areas* represent elevations above 1500 m, where all known summer aestivation sites are located. *ACT* = Australian Capital Territory. Adapted from Green et al. (2021). **Inset, upper left:** The Australian Bogong moth *Agrotis infusa* (Boisduval, 1832). Scale bar = 5 mm. Reproduced with the kind permission of the photographer; Ajay Narendra, Macquarie University, Australia.

Once they have reached the Australian Alps, Bogong moths seek out the cool cracks and crevices of particular granite outcrops dotted across the mountain ridges. Here they spend the summer huddled together in a dormant state known as aestivation (Common, 1952). In autumn, they leave these outcrops and return to their breeding grounds, where they breed, lay their eggs, and die, ending their migratory journey (Green, 2010). The following generation of Bogong moths will return to the same set of summer sites as their ancestors, sites known to have been repeatedly and consistently occupied by Bogong moths for millennia (Keaney et al., 2016; Stephenson et al., 2020). This journey is achieved despite the moth having had no opportunity to learn the location of those sites from the previous generation. The information the moth requires to reliably navigate to its alpine destination during the spring migration must therefore be inherited.

The impressive migration of the Bogong moth, along with its abundance and accessibility, has seen it become an important emerging model for long-distance nocturnal navigation, subject to fruitful and ongoing research which has begun to unravel the sensory and neurobiological mechanisms that the moth uses to successfully migrate (Adden et al., 2020; Dreyer et al., 2018; Warrant et al., 2016). However, there remains an important open question which this study aims to address: How are naïve moths capable of using their navigational toolkit to migrate from broadly distributed breeding grounds to a relatively restricted area with such remarkable species-level site fidelity?

Previously, the degree of linkage between specific regions in the Bogong moth breeding grounds and their migratory destination (i.e. the aestivation sites)—a concept known as ‘migratory connectivity’—was not even known (Gao et al., 2020). Under a scenario where migratory connectivity in the Bogong moth is high, Bogong moths would have a propensity to return after aestivation to the specific region of their breeding grounds where they hatched months earlier. We would then expect genetic structure to develop across the Bogong moth population, as gene flow would necessarily be limited between subpopulations from different breeding regions. Alternatively, if migratory connectivity is low, there would be a high degree of mixing across the Bogong moth population, which would lead to an absence of population structure.

The degree of migratory connectivity present in Bogong moths has important implications for how Bogong migratory direction is inherited. A high-migratory-connectivity scenario would readily facilitate a regime favouring a genetically-determined migratory direction, akin to that seen in migratory songbirds (e.g. Lundberg et al., 2017)—Bogong moths would then simply inherit a fixed migratory direction from their parents. However, in a low-migratory-connectivity scenario, additional information would be required in order for the Bogong moth to successfully complete its migration. This information could come in the form of sensory cues, perhaps relating to the environment in which the moth hatches, or, as in the high-connectivity scenario, it could come in the form of a genetically-heritable fixed spring-migratory direction. Whether or not this inherited migratory direction is correct (for a given breeding ground region) would only be determined once the merciless process of natural selection has acted. This latter possibility may be reasonable, given the high fecundity of Bogong moth females, which can each lay up to 2,000 eggs (Warrant et al., 2016).

In this study, we aim to determine whether Bogong moth migratory direction is genetically heritable, and if so, to infer the putative sources of its heritability. In doing so, we also aim to shed light on the level of migratory connectivity in the Bogong moth. We will proceed by taking advantage of the research opportunities presented by the recently sequenced Bogong moth reference genome (Wallace et al., in prep.). By re-sequencing the whole genomes of 77 Bogong moths collected from locations distributed across their entire breeding grounds and their summer aestivation range, we will search for genetic variation that could explain the variation in their migratory directions.

In addition to progressing our understanding of Bogong moth migration, these population genetics data will also prove useful for understanding Bogong moth ecology more generally. The Bogong moth has the peculiar status of being both endangered (Warrant et al., 2021), and simultaneously considered a pest (Common, 1954; Farrow and McDonald, 1987, see also https://moths.csiro.au/species_taxonomy/agrotis-infusa/). This duality is a direct result of its migratory life history—in summer adult Bogong moths provide essential nutrients and energy to the Australian Alpine ecosystem (Green, 2011), but in winter Bogong cutworms are known to cause damage to crops in their breeding grounds (Common, 1990). Regardless of whether management objectives are conservation or control, an understanding of the structure of the Bogong moth population is fundamentally important for informing strategies to achieve them.

## 2 Methods

### 2.1 Sample material

In the Southern Hemisphere spring months of 2017-2019, living Bogong moths were sampled from various locations across their breeding grounds and spring migratory routes. Moths were attracted to a white sheet suspended in a tree and illuminated by a 1000 W xenon searchlight and/or a LepiLED lamp (Brehm, 2017). Bogong moths were collected by hand using sample jars. Moths were also collected during the summers of 2017-2020 from Bogong aestivation sites in the alpine regions of New South Wales and Victoria. The whole moth samples were fixed in absolute EtOH shortly after collection, and were stored at either room temperature or 4°C. This eliminated logistical issues associated with attempts to keep samples at low temperatures in the field, and proved effective in maintaining the quality of the samples’ DNA. The collection locations are listed in Table 1, and are also included on the map in Fig. 1.

**Table 1:** Sample collection locations. Samples are grouped by two-letter abbreviations (Abbr.). Migratory condition (Cond.) is noted as either spring migrant (SM) or aestivating (A). *n* denotes the number of samples sequenced for each group, and *n*′ denotes the number of samples which passed sequencing quality control.

| Abbr. | Location | Cond. | Lat/Lon | Date | $n$ | $n'$ |
| --- | --- | --- | --- | --- | --- | --- |
| CW | Chetwynd | SM | 37°14'S<br>141°30'E | 10-2018 | 13 | 11 |
| DH | Mt Buller | A | 37°09'S<br>146°26'E | 11-2018 | 12 | 5 |
| GN | Mt Gingera | A | 35°34'S<br>148°46'E | 02-2019 | 5 | 4 |
| HA/HS | Hay | SM | 34°33'S<br>144°52'E | 10-2017 | 9 | 9 |
| KG | Mt Kaputar | SM | 30°17'S<br>150°09'E | 11-2017 | 22 | 19 |
| KZ | Mt Kosciuszko | A | 36°27'S<br>148°16'E | 02-2021 | 8 | 8 |
| SA | Moorak | SM | 37°52'S<br>140°45'E | 10-2018 | 12 | 11 |
| SR | Ken Green Bogong | A | 36°31'S<br>148°15'E | 02-2018 | 10 | 9 |
| WW | Wentworth | SM | 34°05'S<br>141°54'E | 10-2018 | 1 | 1 |

### 2.2 DNA extraction

To reduce the chance of contamination, brain and thoracic muscle tissue were dissected from the moth samples and stored at -80°C prior to DNA extraction. DNA was extracted from the brain and muscle tissue separately using a *Quick-DNA™ MagBead Plus kit* (Zymo Research, 2017) and quantified using a *Qubit™ dsDNA HS Assay Kit* on a *Qubit™ 3 Fluorometer* (Thermo Fisher Scientific, 2017) using the manufacturers’ respective protocols. Results from the first few extractions indicated that DNA yield was significantly higher from the thoracic muscle tissue when compared with the brain tissue, so for the remainder of the samples DNA was extracted from the muscle tissue only.

### 2.3 Genomic DNA sequencing

A total of 92 Bogong moth samples were selected for sequencing, ensuring sufficient DNA yields for each selected sample, as well as adequate coverage of their geographic range in both the breeding grounds and their mountainous summer aestivation range. Sequencing libraries were prepared using the *Illumina® DNA PCR-Free Prep, Tagmentation* with the *IDT® for Illumina® DNA/RNA UD Indexes Set A, Tagmentation* (Illumina, 2020) using the manufacturer’s protocol, and a unique index was used for each sample.

The DNA content of the libraries were quantified using a *Qubit™ ssDNA Assay Kit* on a *Qubit™ 3 Fluorometer* (Thermo Fisher Scientific, 2017), then pooled in equal DNA proportions. To ensure library quality, the pooled library was first sequenced on the Illumina MiSeq platform with a target of 1 million reads, and quality control checks were performed. Based on the read counts from the MiSeq run, the libraries were re-pooled and sequenced on the Illumina NovaSeq platform, for a target of 10-20x average coverage per sample.

### 2.4 Sequencing quality control

For both the MiSeq and NovaSeq runs, quality control was performed using a custom pipeline written in Snakemake (Mölder et al., 2021). Read trimming was performed using Cutadapt (Martin, 2011). To assess contamination levels, trimmed reads were classified using Kraken 2 (Wood et al., 2019) with the National Center for Biotechnology Information (NCBI) non-redundant nucleotide (nt) database, and were aligned to the Bogong reference genome (Wallace et al., in prep.) using BWA-MEM 2 (Vasimuddin et al., 2019). General quality control statistics were obtained using FastQC, SAMtools (Li et al., 2009), and Picard tools “collectinsertsizemetrics.” Summary reports were generated from the output of the above software using MultiQC (Ewels et al., 2016).

### 2.5 Variant calling

Sequence variant (SNP/INDEL) calling was performed using the data from both the MiSeq and NovaSeq runs, using a slightly modified version of the variant-calling pipeline, Grenepipe (Czech and Exposito-Alonso, 2021). Read trimming was performed using Trimmomatic (Bolger et al., 2014), mapping was performed using BWA-MEM 2 (Vasimuddin et al., 2019), and variant calling was performed using GATK haplotypecaller (McKenna et al., 2010).

Exon drop-out variants were identified by assessing read coverage of exons in the reference genome annotation, based on the mapping generated in the sequence variant calling step. Exons which had no reads mapped to them from a given sample were inferred to be missing from that sample’s genome. To reduce incidence of false discoveries of exon drop-outs, only samples with an average read-depth across all exons above a certain threshold were considered. From our analyses, an average read-depth of 10x is sufficient to bring the false-discovery rate for most exons to a vanishingly small value, well below 10^−5^, however lower average read-depths perform considerably worse, so a threshold of 10x was used (see Appendix A.1 for details of the analysis).

Transcript drop-out variants were identified in a similar fashion to exons. For further analyses, only a single representative transcript for each unique pattern of presence across the samples within each gene was used. That is, if a transcript shared identical presence/absence information with another transcript from the same gene which had already been included, then the former would be discarded from subsequent analyses. In this way, transcript drop-outs approximate gene drop-outs.

### 2.6 Population structure analysis

Population structure was assessed based on the sequence variant data using Structure_threader (Pina-Martins et al., 2017) and fastStructure (Raj et al., 2014) for *k* values ranging from 1–12. Principal component analysis was also performed on the sequence variant data. Clustering based on the hamming distance from sequence variant data and transcript drop-out variant data was also done using the neighbour-joining algorithm.

### 2.7 Genome-wide association study of migratory direction

To determine if any variants are associated with migratory direction, we performed a genome-wide association study (GWAS) on spring migratory direction. To perform the analysis, spring migrants were classified as either western (Chetwynd, Hay, Moorak, and Wentworth samples) or northern (Mt Kaputar samples). Summer aestivating moths (Mt Gingera, Mt Kosciuszko, and Ken Green Bogong samples) were not included in the GWAS, as it is not known from which direction they migrated in the preceding spring. Association analysis was performed with univariate linear mixed models using GEMMA (Zhou and Stephens, 2012). P-values were calculated using a likelihood ratio test and a significance threshold of 0.05 with Bonferroni correction was applied. To limit the study to variants which could have large effect sizes, we set a strict minimum minor allele frequency threshold of 0.3.

## 3 Results

### 3.1 DNA sequencing and quality control

Of the 92 Bogong moth samples selected for sequencing, 77 passed all quality control steps (refer to Table 1 for the locations of the successful samples). The failed samples were excluded due to high levels of contamination or sample drop-out during sequencing. We believe the sample drop-outs were caused by a technical error which resulted in one column (8 samples) of the library preparation failing. Summaries of the results of quality control on the main sequencing run are shown in Appendix A.2 Fig. A.2.

From the Kraken 2 read classification analysis, we identified three species of bacteria which occurred in large quantities or in multiple samples. These were *Providencia rettgeri, Alcaligenes faecalis*, and *Serratia marcescens*. Interestingly, *P. rettgeri* is known to be pathogenic to insects, and is carried by insect-parasitic nematodes of the genus *Heterorhabditis* (Jackson et al., 1995). Adult Bogong moths are known to be parasitised by two species of mermithid nematodes (Common, 1954; Welch, 1963), although it is not known if these are also carriers of *P. rettgeri*.

### 3.2 Variant calling

A total of 186,622,107 sequence variants were discovered, representing approximately one variant for every 3.18 bases in the reference genome, exceeding the notable levels of variation recently reported in the diamondback moth *Plutella xylostella*, which has approximately one variant every 6 bases (You et al., 2020). Of the discovered sequence variants, 153,363,045 were SNPs and 33,259,062 were INDELs. Approximately half of the sequence variants were singletons (74,336,046 SNPs and 19,300,939 INDELs occurred in only a single sample).

A total of 11,252 exon drop-out variants and 846 transcript drop-out variants were discovered. Of the transcript drop-out variants, 258 were singletons. We performed gene ontology enrichment analysis on these 846 transcripts and found a small number of terms which were significantly enriched (greater number of occurrences than would be expected by chance), and a single term which was significantly purified (fewer occurrences than would be expected by chance), when compared to the rest of the Bogong moth gene set (*p <* 0.05, Fisher’s exact test, Bonferroni corrected). These terms are presented in Table 2.

**Table 2:** Gene ontology terms which were significantly enriched or purified in the set of transcripts which were not present in every Bogong moth sample sequenced (*p <* 0.05, Fisher’s exact test, Bonferroni corrected).

| Term | Name | $p$ |
| --- | --- | --- |
| **Enriched terms** |  |  |
| *Biological process* |  |  |
| GO:0015074 | DNA integration | $3.58 \times 10^{-26}$ |
| GO:0006915 | apoptotic process | $1.73 \times 10^{-6}$ |
| GO:0006313 | transposition, DNA-mediated | $9.28 \times 10^{-9}$ |
| GO:0006310 | DNA recombination | $2.81 \times 10^{-10}$ |
| *Cellular component* |  |  |
| GO:0000786 | nucleosome | $4.73 \times 10^{-25}$ |
| *Molecular function* |  |  |
| GO:0004803 | transposase activity | $2.06 \times 10^{-9}$ |
| GO:0046982 | protein heterodimerization activity | $6.94 \times 10^{-30}$ |
| GO:0003676 | nucleic acid binding | $3.28 \times 10^{-20}$ |
| GO:0003677 | DNA binding | $1.74 \times 10^{-19}$ |
| **Purified terms** |  |  |
| *Molecular function* |  |  |
| GO:0005515 | protein binding | $6.25 \times 10^{-11}$ |

### 3.3 Population structure

No evidence of population substructure was found based on sequence variant (Fig. 2a–e) or transcript drop-out variant (Fig. 2f) data. Structure analysis showed that the data are best explained by a single panmictic population (Fig. 2a–c). This was supported by principal component analysis (Fig. 2d) which failed to produce any meaningful sub-population clusters. The neighbour-joining algorithm applied to sample-pair-wise hamming distances of sequence or transcript drop-out variants also failed to indicate any meaningful population structure (Fig. 2e–f).

**Figure 2:**
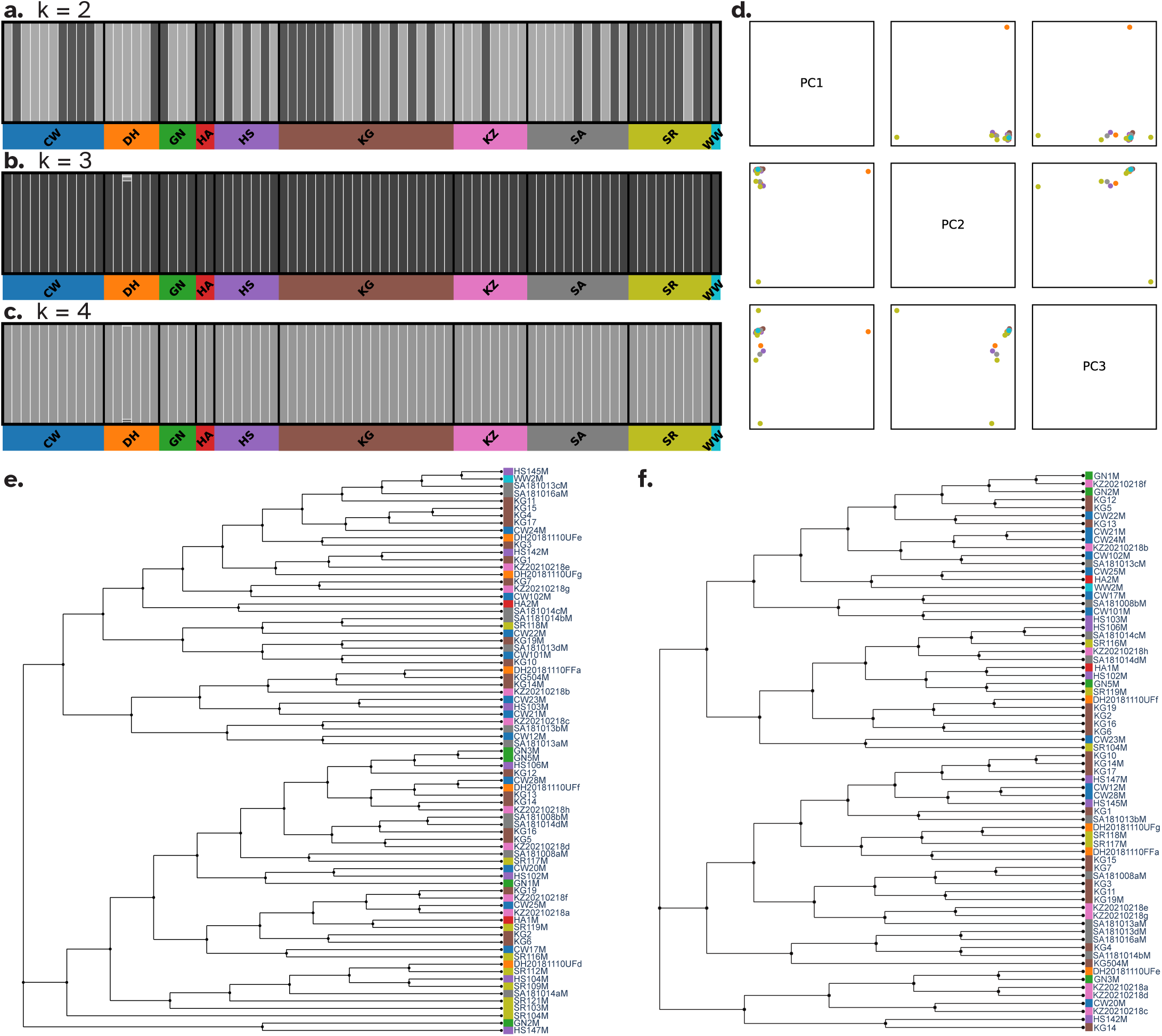
No evidence of population structure was detected in variant data from the largest 31 scaffolds of the Bogong moth reference genome, amongst samples collected from across the Bogong moth breeding grounds and summer aestivation range. **a–c**. Structure plots with values of *k* set to 2, 3, and 4 (*k* values of 1,5–12 were also tested, but have been omitted for brevity) produced using fastStructure (Raj et al., 2014). Two-letter geographical location abbreviations are defined in Table 1 and shown in Fig. 1. The data were found to be best explained using *k* = 1 (i.e. a single panmictic population, although the plot for *k* = 1 is not shown as it is uninformative, since *k* = 1 is the trivial case). **d**. The first three principal components of sequence variants. Samples are coloured by location, as in a–c. We did not observe any clear clustering into Bogong moth subpopulations. **e–f**. Neighbour-joining dendrograms from hamming distance of sequence variant (**e**) and transcript drop-out variant (**f**) data. Samples are coloured by location, as in a–c. No evidence of population stratification was detected in either dataset.

### 3.4 Genome-wide association study

Four sequence variants were found to be significantly associated with migratory direction to the Bonferroni-corrected significance level of *p <* 1.13 *×* 10^−7^ (Fig. 3). We refer to these variants as AiSNP_01, AiSNP_02, AiSNP_03, and AiSNP_04. Summaries of the genotype proportions of these significantly-associated loci in each of the sample collection locations are shown in Fig. 4. Summaries of the genomic context of each variant, with reference to functional information from orthologs studied in other species, are provided in Table 3.

**Table 3:** Summary of loci which are significantly correlated with spring migratory direction.

| ID/ Scaffold:<br>Coordinate | Description |
| --- | --- |
| AiSNP_01/<br>HiC_scaffold_9:<br>16593335 | $p = 6.87 \times 10^{-10}$. Located in the 3' exon of an uncharacterised gene, which is expressed in eye and brain tissue in the Bogong moth. The SNP maps to the third position of a tyrosine codon, and is a synonymous substitution (TAT → TAC). |
| AiSNP_02/<br>HiC_scaffold_12:<br>4434220 | $p = 8.06 \times 10^{-8}$. Appears to be in a non-coding region, with the closest gene being approximately 10 kb away. |
| AiSNP_03/<br>HiC_scaffold_15:<br>11378716 | $p = 1.73 \times 10^{-8}$. Located in an intron, towards the 5' end of the OUTD7B gene (OTU domain-containing protein 7B), which is involved in ubiquitination, and ultimately immune regulation (Hu et al., 2016, 2013), DNA repair, and the EGFR/MAPK (Lei et al., 2019) and mTORC2 (Wang et al., 2017) pathways. |
| AiSNP_04/<br>HiC_scaffold_25:<br>12095901 | $p = 2.14 \times 10^{-7}$. Located in the coding region of the NFXL1 gene (NF-X1-type zinc finger protein NFXL1). NFXL1 is thought to mediate DNA binding and function as an E3 ubiquitin ligase in plants (Müssig et al., 2010), and has been shown to be expressed in human embryonic stem cells, but down-regulated during differentiation to myelinated oligodendrocytes (Chaerkady et al., 2011). Variants in the human NFXL1 gene have been associated with risk of diagnosis with Specific Language Impairment during childhood (Villanueva et al., 2015). The SNP maps to the third position of a glutamic acid codon, and is a synonymous substitution (GAA → GAG). |

**Figure 3:**
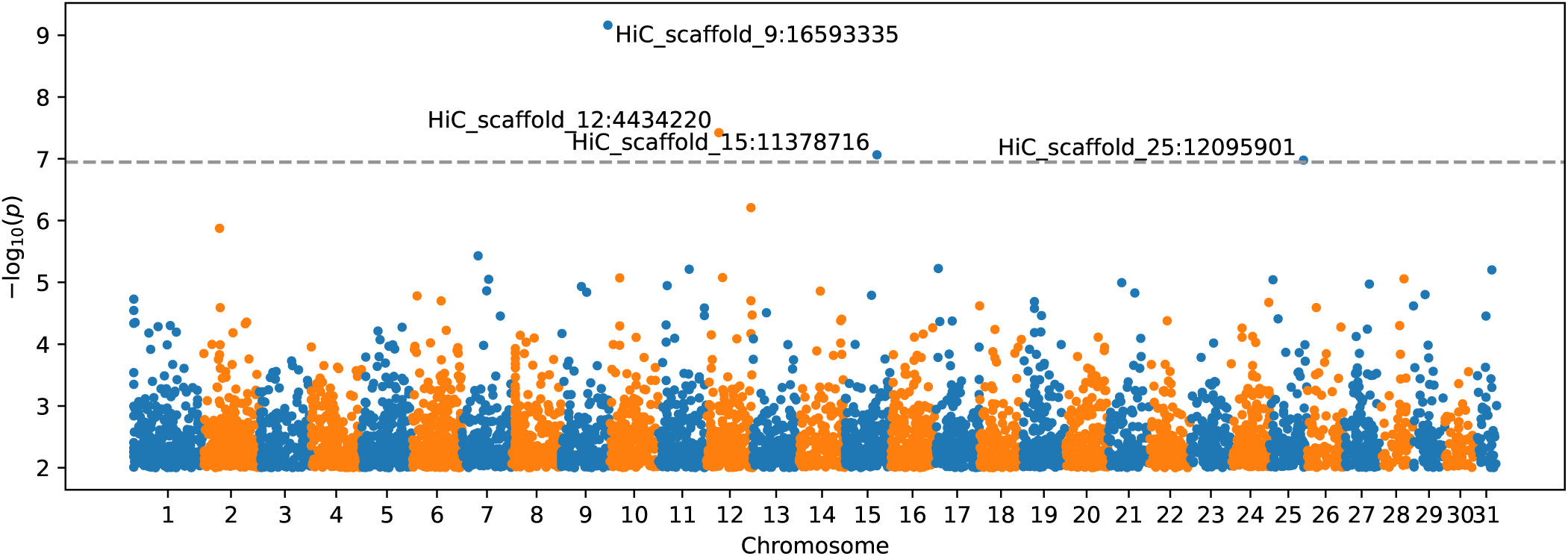
Manhattan plot of sequence variant associations with spring migration orientation as a categorical variable. Spring migrants were categorised as either “south-flying” or “east-flying,” depending on the location where they were caught (see Section 2). *Dashed line* indicates Bonferroni p-value threshold of 1.13 × 10^−7^. The four significantly associated variants are labelled. Loci with p-values greater than 0.01 are omitted. Points are coloured according to which scaffold (chromosome) they appear in, with odd-numbered scaffolds coloured blue and even-numbered scaffolds coloured orange.

**Figure 4:**
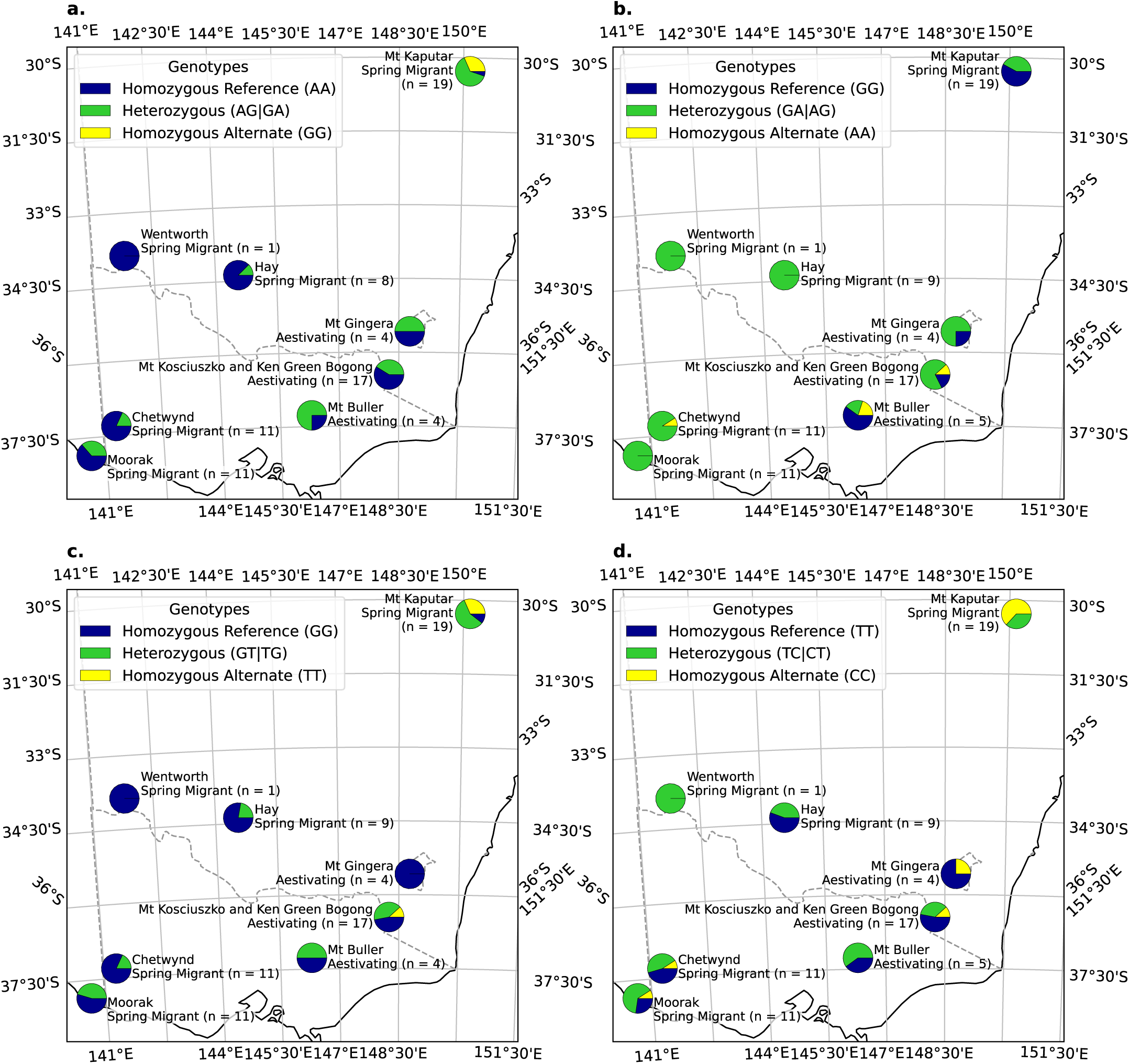
Genotype proportions for sequence variants significantly associated (*p <* 0.05, Bonferroni-adjusted *p <* 1.13 × 10^−7^) with spring migration orientation according to geographic location across the Bogong moth range. Plotted using Azimuthal Equidistant map projection, observer centred on Mt Kosciuszko. **a**. AiSNP_01 (HiC_scaffold_9:16593335, *p* = 6.87 *×* 10^−10^). **b**. AiSNP_02 (HiC_scaffold_12:4434220, *p* = 8.06 *×* 10^−8^). **c**. AiSNP_03 (HiC_scaffold_15:11378716, *p* = 1.73 *×* 10^−8^). **d**. AiSNP_04 (HiC_scaffold_25:12095901, *p* = 2.14 *×* 10^−7^).

### 3.5 Regions under selection

Tajima’s D (Tajima, 1989) is a test statistic frequently used to assess the mode of selection acting on nucleotide sequences. It evaluates the null hypothesis that a sequence of DNA has evolved under neutral selection by comparing estimates of nucleotide diversity based on the number of polymorphic sites with estimates based on the allele frequencies of those sites. Values of Tajima’s D which deviate negatively from 0 indicate an over-abundance of rare variants, with respect to the expectation under neutral selection, suggesting a recent selective sweep or population expansion. Values of Tajima’s D which deviate from 0 in the positive direction indicate an over-abundance of common variants, with respect to the neutral expectation, suggesting the presence of balancing selection or a population contraction.

We calculated Tajima’s D in 10 kb windows across the Bogong moth genome using VCF-kit 0.2.9 (https://github.com/andersenlab/VCF-kit). A marked depression of Tajima’s D was observed across the genome (Fig. 5), implying an over-abundance of rare variants.

**Figure 5:**
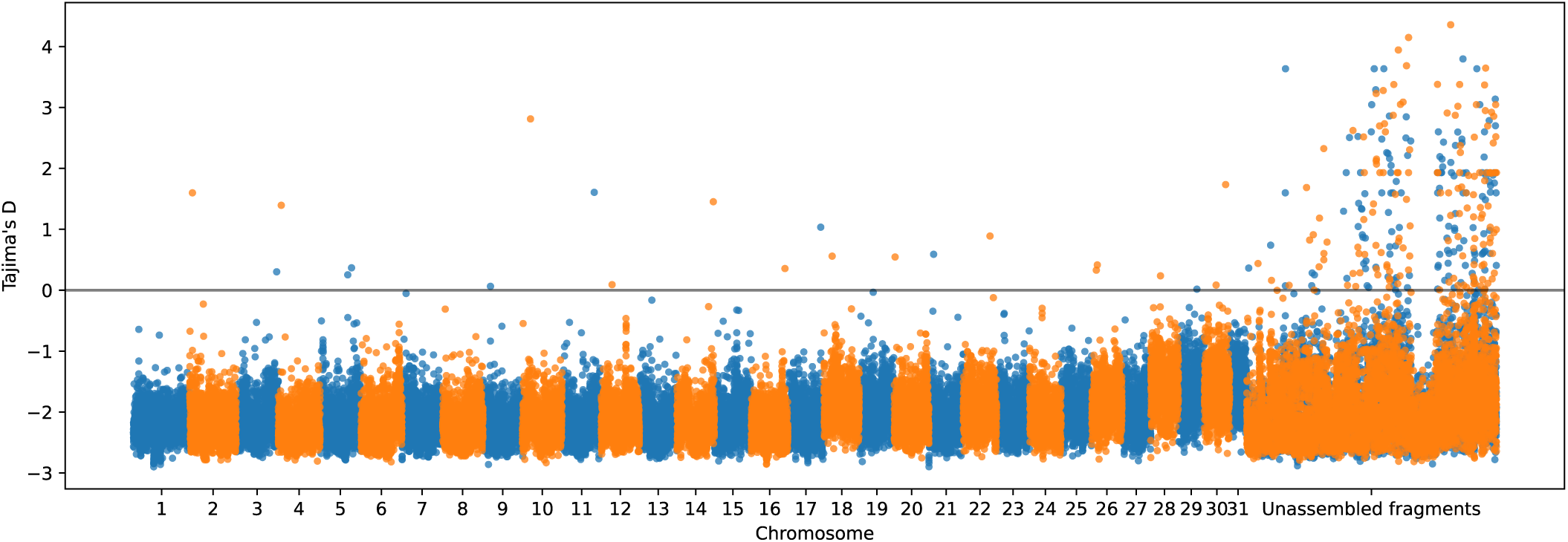
Genome-wide scan of Tajima’s D (Tajima, 1989), calculated in non-overlapping 10 kb bins. Colouring convention follows that of Fig. 3.

We identified genes which are likely to have recently been acted on by balancing selection or positive selection, by selecting genes which overlapped with the top or bottom 1% of the Tajima’s D bins, respectively. A full list of these genes is presented in Appendix A.3. A total of 376 genes were identified in the top 1% of the Tajima’s D bins (full list: Table A.1), and 256 genes were in the bottom 1% (full list: Table A.2). To determine if selection was acting on particular biological processes or molecular functions, we performed a gene ontology enrichment analysis on both groups of identified genes and found multiple significantly enriched terms in each (*p <* 0.05, Fisher’s exact test, Bonferroni corrected). These enriched terms are presented in Table 4.

**Table 4:**
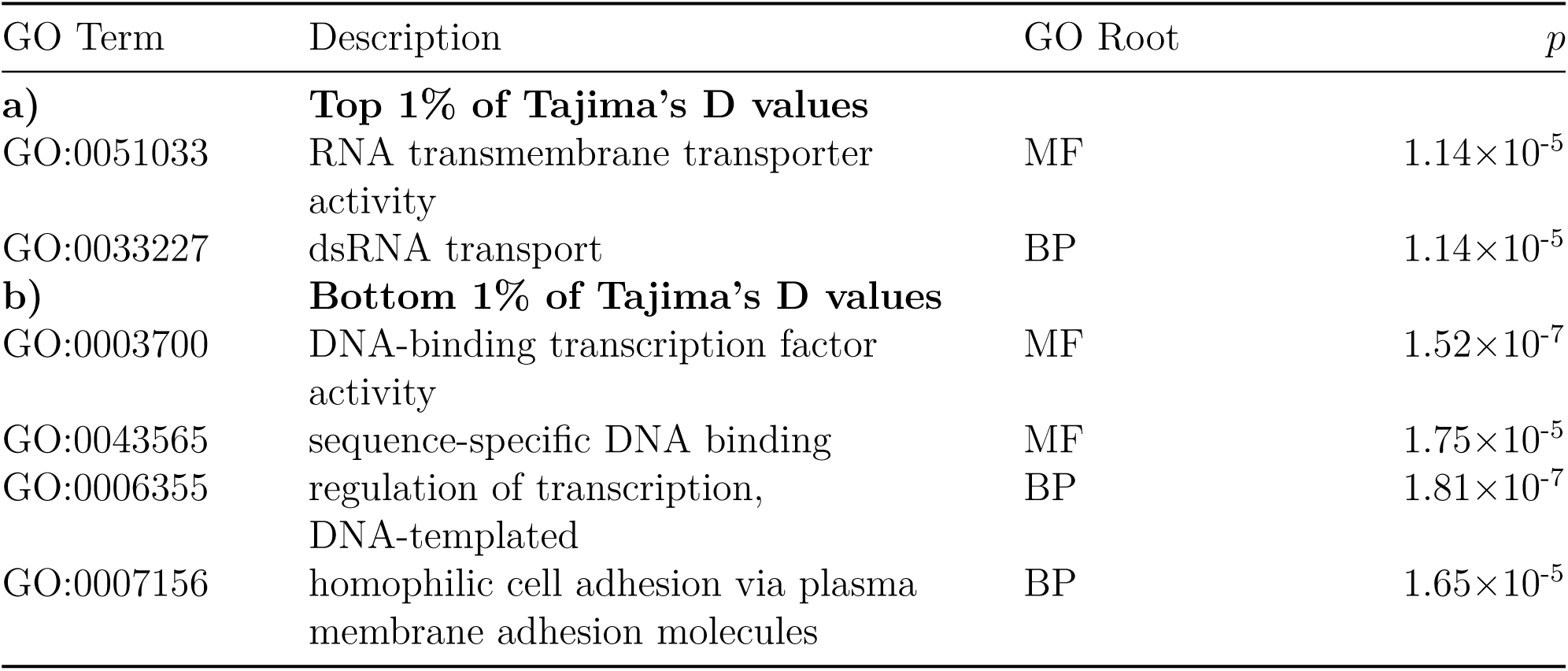
Enriched gene ontology terms in genes co-located with the top (**a**) and bottom (**b**) 1% of Tajima’s D values, calculated in 10 kb bins across the first 31 scaffolds in the Bogong moth genome. Terms shown were found to be significantly enriched (*p <* 0.05, Fisher’s exact test, Bonferroni corrected). GO Root terms shown are Molecular Function (MF) and Biological Process (BP).

| GO Term | Description | GO Root | $p$ |
| --- | --- | --- | --- |
| **a)** | **Top 1% of Tajima’s D values** |  |  |
| GO:0051033 | RNA transmembrane transporter activity | MF | $1.14 \times 10^{-5}$ |
| GO:0033227 | dsRNA transport | BP | $1.14 \times 10^{-5}$ |
| **b)** | **Bottom 1% of Tajima’s D values** |  |  |
| GO:0003700 | DNA-binding transcription factor activity | MF | $1.52 \times 10^{-7}$ |
| GO:0043565 | sequence-specific DNA binding | MF | $1.75 \times 10^{-5}$ |
| GO:0006355 | regulation of transcription, DNA-templated | BP | $1.81 \times 10^{-7}$ |
| GO:0007156 | homophilic cell adhesion via plasma membrane adhesion molecules | BP | $1.65 \times 10^{-5}$ |

## 4 Discussion

Our analyses show that the Bogong moth population contains a large amount of (mostly rare) variation, and that this variation does not correspond to the geographic distribution of its winter breeding grounds, nor its summer range. In fact, the variation appears to be entirely unstructured in the population. Therefore, we conclude Bogong moths form a single panmictic population, a result which agrees with previous attempts to characterise Bogong moth population structure using SNP array (Peter Kriesner, personal communication) and RNA-Seq data (Wallace et al., in prep.). A similar result has recently been reported in the Bogong moth’s diurnal Northern Hemisphere migratory counterpart, the monarch butterfly *Danaus plexippus* (Talla et al., 2020).

Panmixia of the Bogong moth population requires high levels of gene flow between the far reaches of their breeding grounds. This suggests that individual Bogong moths do not, in general, return to the specific region in the breeding grounds from which they originated once they have completed their high-altitude aestivation—at least lineages of moths do not consistently return to a specific region over multiple generations. That is, it is likely that Bogong moths have low migratory connectivity.

Despite the suitability of panmixia as a model of Bogong moth population structure, we were able to discover a small number of variants which were significantly associated with geographic location, and, by inference, migratory direction. Three of these variants occur within genes, however the functional consequences of the variants are not immediately obvious. Two of the variants cause synonymous substitutions and one is located in an intron, so putative function is likely to be conferred through some regulatory process, rather than a modification of a gene product *per se*. Nevertheless, further investigation into the functional significance of each of these variants is warranted.

We are left with a curious combination of conclusions. On the one hand, it seems likely that Bogong moth population and migratory connectivity is low, meaning the moths are not predestined to return to the specific region they hatched in order to breed. On the other hand, there are some some genetic variants associated with geographic location in the breeding grounds. An obvious question arises—how are genetic associations with geographic location established and/or maintained in a panmictic population? An enticing possible explanation is that these location-associated variants are subject to location-dependent selection pressure, which would reduce the capacity of gene flow to eliminate the location-dependent variation. Either a geographically-imprecise return migration of the Bogong moths to their breeding grounds or a degree of mixing and mating prior to commencing the return migration^1^ could then facilitate enough gene flow to remove signs of population structure across the broader genome.

An imprecise return migration could be considered adaptive, as it would enable rapid re-colonisation of breeding areas rendered temporarily unfavourable by climatic events such as drought (Farrow and McDonald, 1987), or even untimely or excessive rainfall, which can affect noctuid pupal survival (Murray and Zalucki, 1990a, 1990b; Sims, 2008). Indeed, rapid re-colonisations by Bogong moths (and other noctuid moths) do occur (Farrow and McDonald, 1987). The offspring of these colonists would benefit from some degree of flexibility in their putative inherited migratory direction, as they migrate from a different starting area from that of their parents, who migrated the previous year.

As with any association study, the associations we have discovered in our genome-wide scan for location-associated genetic variants are simply correlations, and causality needs to be confirmed with reverse genetics experiments. Such experiments should include direct measurement of the phenotype of interest, namely, migratory direction. This could be achieved using a Frost-Mouritsen flight simulator, which has an established use for studying Bogong moth migration (Dreyer et al., 2021). This would enable the experiment to disentangle potentially confounded phenotypic and behavioural responses, such as timing of migration and migratory direction. Such confounding factors may complicate the present study, owing to the variation in collection time for samples from different areas (Table 1). However, for the purposes of our aims, they are not of great concern, as genetic correlations to migratory timing are interesting in their own right, and would still require reverse-genetic confirmation.

Reverse genetics experiments that measure a behavioural phenotype are necessarily laborious, and therefore only tractable for studying a small number of genes. Equipped with the results from this large-scale sequencing effort, we now know where to look for genes which are putatively involved in controlling long-distance navigation, opening the door to previously intractable experiments which could shed light on the fascinating phenomenon of directed animal migration.

The whole-genome scan for signals of selection yielded evidence of a marked over-abundance of rare variants, with respect to our expectation under the null hypothesis that the genome sequence evolved under neutral selection and fixed population size. This indicates that the Bogong moth population has likely recently recovered from a past genetic bottleneck, resulting in variation being dominated by mutations rather than genetic-drift-mediated partial fixation. It is known that Bogong moth population size can vary dramatically from year to year (Green et al., 2021), although it is thought that this has been underpinned by a slow downward trend over the last half-century (*ibid*.).

Interestingly, regions of the genome with high Tajima’s D values appear to be preferentially co-located with genes involved in RNA transmembrane transporter activity (Table 4a). High values of Tajima’s D are often interpreted as evidence of balancing selection, or recent population contraction. The latter seems unlikely because most of the genome has strongly negative Tajima’s D values (Fig. 5). That leaves us with the conclusion that balancing selection is acting on these regions to increase their diversity in the population. RNA transport is important for regulation of gene expression, which is a fundamental process, so changes in how it operates could have far-reaching consequences for the organism. It is therefore not hard to imagine that this type of selection acting on these regions could have important ecological implications.

On the other hand, genomic regions with low Tajima’s D values appear to be preferentially co-located with genes involved in sequence-specific transcription regulation, as well as homophilic cell adhesion (Table 4b). Low values of Tajima’s D are often interpreted as evidence of a recent selective sweep (i.e. positive selection) or recent population expansion. Since we are looking only at regions which have the bottom 1% of Tajima’s D values, we can be fairly confident that selective sweeps are good explanations for the low values near these genes, albeit in a background of population expansion. The enrichment of positive selection acting on sequence-specific transcription regulation is particularly interesting in the context of the Bogong moth, as these are the type of potential molecular events that can bring about important behavioural changes already seen in other organisms (Rittschof et al., 2014).

It is reasonable to assume that selection acts strongly on migration and migratory direction in the Bogong moth. Directed migration in lepidoptera is known to be remarkably fragile (Tenger-Trolander et al., 2019), and would probably disappear quickly without the continual action of selection. It is therefore plausible that migratory direction itself is the phenotype under location-dependent selection, leading to genotypic selection on the genomic regions and the location-dependent variants we identified. Perhaps the most promising location-dependent variant is AiSNP_03, which is located in an intron towards the 5’ end of the OUTD7B gene. OUTD7B is involved in immune regulation (Hu et al., 2016, 2013), and the EGFR/MAPK (Lei et al., 2019) and mTORC2 (Wang et al., 2017) pathways, which are central to regulatory networks in the cell, and act in a context-dependent manner. One could speculate that AiSNP_03 interferes with an intronic enhancer leading to changes in a regulatory module, with broader implications for gene expression.

A common theme has emerged from our analyses of 77 Bogong moth genomes: genetic regulation. Animal behavioural properties (such as navigational skills) are largely emergent and depend on selective events at molecular or physiological levels. It is therefore conceivable that subtle changes to regulation—mediated by modulation of important biochemical pathways such as EGFR/MAPK and mTORC2—could have profound implications for behavioural control. Indeed, regulatory networks must be extremely important for differentiating between the migration and aestivation phases in the Bogong moth life cycle, as deep differences in the transcriptional profiles of moths in these two behavioural states have been observed in sensory and brain tissues (Wallace et al., in prep.).

Bogong moths’ remarkable behaviour is a distributed property of many brain networks controlling sensory organs, muscles, metabolic flux, etc. that respond to both external and internal cues. For example, the encoding of either north/south or east/west directionality could be described as an inherited “value,” selected through evolutionary and somatic processes (Friston et al., 1994). The Bogong moths’ preference for cooler areas have likely strengthened certain neuronal connections and created this value (e.g. during spring, south is better than west for the Bogong moth, much like dark is better than light for many animals). When this value is realised, these connections are reinforced and dominate amongst numerous neuronal connections (Sporns et al., 2000). When external or internal conditions change—say, the temperature increases, or the moth’s brain reaches a certain stage of maturation—these neurons fire together, driving the moth’s desire to move towards cooler regions, which they achieve through integration of other navigationally-informative cues, (e.g. landmarks and the geomagnetic field, Dreyer et al., 2018; the night sky, Adden, 2020).

To implement such a value system, the Bogong moth must inherit the capacity to sense relevant thresholds in the cues which inform its migration (e.g. temperature, day length, celestial cues, geomagnetism, etc.). For example, there must be a set of thresholds that either allow or prevent the onset of migration, as a premature or delayed migration could have severe consequences. Similarly, the Bogong moth must have a way of measuring compass directions, including—but not limited to—detecting the geomagnetic field (Dreyer et al., 2018), possibly using a sensor similar to cryptochrome 4, which is thought to be the magnetoreceptor in night-migratory songbirds (Xu et al., 2021). Such capacities are established via a developmental program that has evolved some Bogong moth-specific features, but overall, cannot be hugely different from its close non-migratory—or non-directionally-migratory—relatives.

To become heritable, a behavioural novelty must be reflected in the genome. For example, the migratory direction-associated SNPs we discovered in this study could represent variations which provide a selective advantage for the migratory behaviour of the Bogong moth. SNPs in non-coding (intronic or intergenic) regions are of special interest because they are more likely to affect gene expression. We have discussed the intronic AiSNP_03 variant above, and another of the SNPs we discovered is also in a non-coding region (AiSNP_02; Table 3). The remaining two SNPs are in the coding region of genes expressed in eye and brain tissue, and one of which (AiSNP_04) is likely to function as a transcription factor. Of course, there are many more SNPs that may not show in statistical tests, but could still influence gene regulation.

An important mechanism for far-reaching regulatory control is epigenomic modification, mediated through the covalent bonding of a methyl group to a base (typically a cytosine). Even small genomic changes often involving epigenomic modifiers have been shown to have a massive effect on brain development and function (reviewed by O’Donnell and Meaney, 2020). Importantly, there are a number of ways that epigenomic modifications can be inherited. First, there appears to be a lot of heterogeneity in the Bogong moth genome (which is, by definition, heritable) on which selection could act at the epigenomic level. For instance, in the honeybee genome, there are over 220,000 SNPs that potentially could change the number of methylated sites, either by changing a CpG dinucleotide (the typical target of DNA methylation) to another (e.g. ApG), or by creating a new target (e.g. by changing a TpG to CpG) (Wedd et al., 2016). It is likely that there are even more such SNPs in the Bogong moth genome. Second, we also now understand how acquired (epigenetic) features could be transferred to the next generation, for example, by microRNAs in sperm (reviewed by Chen et al., 2016). Therefore, there are molecular mechanisms by which the entire gene-regulatory network topology can be modified and rewired to generate novel gene expression patterns, and these can likely be passed from one generation to another.

Thus, our analysis of genetic variation in the Bogong moth is yet another example of how epigenetic mechanisms, bound by genetic constraints, are prime drivers of brain plasticity arising from both developmental and experience-dependent events. Furthermore, our results expose a multitude of interesting and exciting avenues of further research into the molecular basis of insect migration, and help establish the Bogong moth as an illuminating emerging model, not only for the study of nocturnal migration and navigation, but also for the study of the fundamental biomolecular processes that contribute to complex animal behaviour.

## 5 Author contributions

JRAW, EJW, and RM conceived the project and designed the experiment. JRAW made the majority of sample collections, coordinated the remainder of the collections, performed the lab work, analysed the data, and wrote the draft version of the manuscript. All authors interpreted the results, contributed text, and edited the manuscript.

## 6 Acknowledgements

EJW and JRAW are grateful for funding from the European Research Council (Advanced Grant No. 741298 to EJW), and the Royal Physiographic Society of Lund (to JRAW). JRAW is thankful for the support of an Australian Government Research Training Program (RTP) Scholarship. EJW holds Scientific Permits for collection and experimental manipulations of Bogong moths in several alpine national parks and nature reserves (NSW: Permit SL100806; Vic: Permit 10008966). The computations and data handling were enabled in part by resources in projects SNIC 2018/8-364 and SNIC 2021/23-74 provided to EJW and JRAW by the Swedish National Infrastructure for Computing (SNIC) at UPPMAX, partially funded by the Swedish Research Council through grant agreement no. 2018-05973.

We thank Ken Green, Dean Heinze, David Szakal, Adrien Lefèvre and Benjamin Mathews-Hunter for assistance with making collections in the field, Niccy Aitken and Justin Borevitz for laboratory resources and guidance, and Jochen Zeil, Robert Kucharski, Gavin Huttley and Andy Bachler for useful comments relating to the analyses.

## A Appendix

### A.1 Expected false discovery rate of missing genomic features from shotgun sequencing

Consider an experiment with the aim of determining whether a particular feature present in a species’ reference genome exists in the genome of an individual sample of that species. In this experiment, the only data available are the reference genome, an annotation of the feature of interest, and whole genome shotgun sequencing reads from the sample. After mapping the reads to the reference, we decide if the feature is present by checking whether there are any reads which map to it. Shotgun sequencing reads are by definition randomly sampled from the genome of the sample. Therefore, the above-mentioned protocol could, by chance, lead to a false inference that the feature is missing from the sample genome. We wish to know how likely these false discoveries are. Equivalently, we wish to know the probability of shotgun sequencing reads missing an extant genomic feature by chance.

For the sake of tractability, we start with a number of assumptions:

1. Read sampling is unbiased.
2. Mapping is perfect.
3. Chromosomes do not have ends (e.g. they are circular. This is roughly equivalent to the feature of interest being far from the ends of a linear chromosome).
4. The feature is a single contiguous genomic region (e.g. a prokaryotic gene or eukaryotic exon).
5. The feature is considered present if any aligned read overlaps the feature by at least *m* nucleotide bases.

In reality, we expect 1 and 2 to hold (at least approximately) for genomic regions which are not redundant or highly repetitive. However, the validity of 2 may also be affected by the particular mapping software used, particularly in the context of sequences which exhibit substantial divergence between the sample and the reference. In prokaryotes, 3 holds, and it also approximately holds in eukaryotes with long chromosomes.

Let the length of the reference genome be *G*, the number of reads *n*, the read-length *k*, and the length of the genomic feature *l*. Then, given assumptions 1 and 3, the probability *p*, that a particular read overlaps the feature by *m* bases is given by

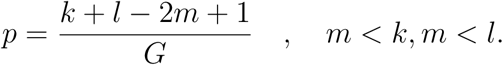

Therefore, the probability, which we will call *Q*, that *n* reads miss the region is given by

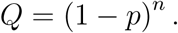

Note that the average sequencing depth *D* is given by

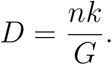

Combining the above equalities gives

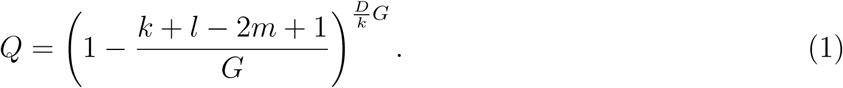

In practice, and particularly when working with eukaryotic genomes, *G* is very large. It is therefore reasonable to use the asymptotic approximation of eq. 1 as *G → ∞*. We begin with a change of variables, letting *a* = *k* + *l −* 2*m* + 1, and 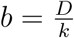. Then

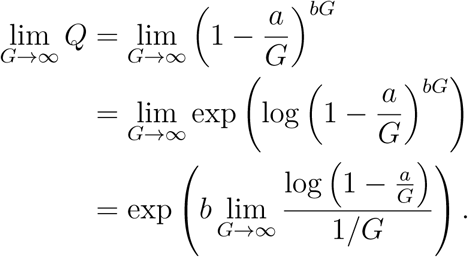

Applying l’Hôpital’s rule, we obtain

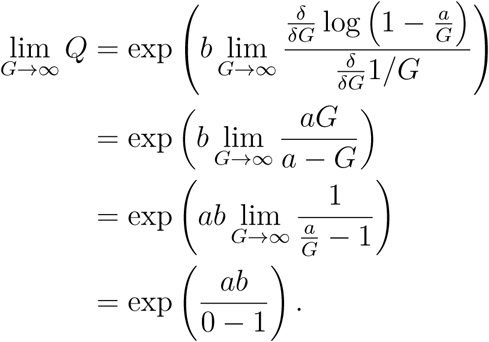

Substituting in the values for *a* and *b* gives us the asymptotic estimate of the probability of missing a feature by chance alone, given the above-mentioned assumptions. Namely,

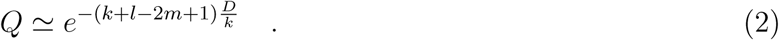

Fig. A.1 shows *Q* plotted against *l* for *k* = 150, *m* = 1, and various values of *D*.

**Figure A.1:**
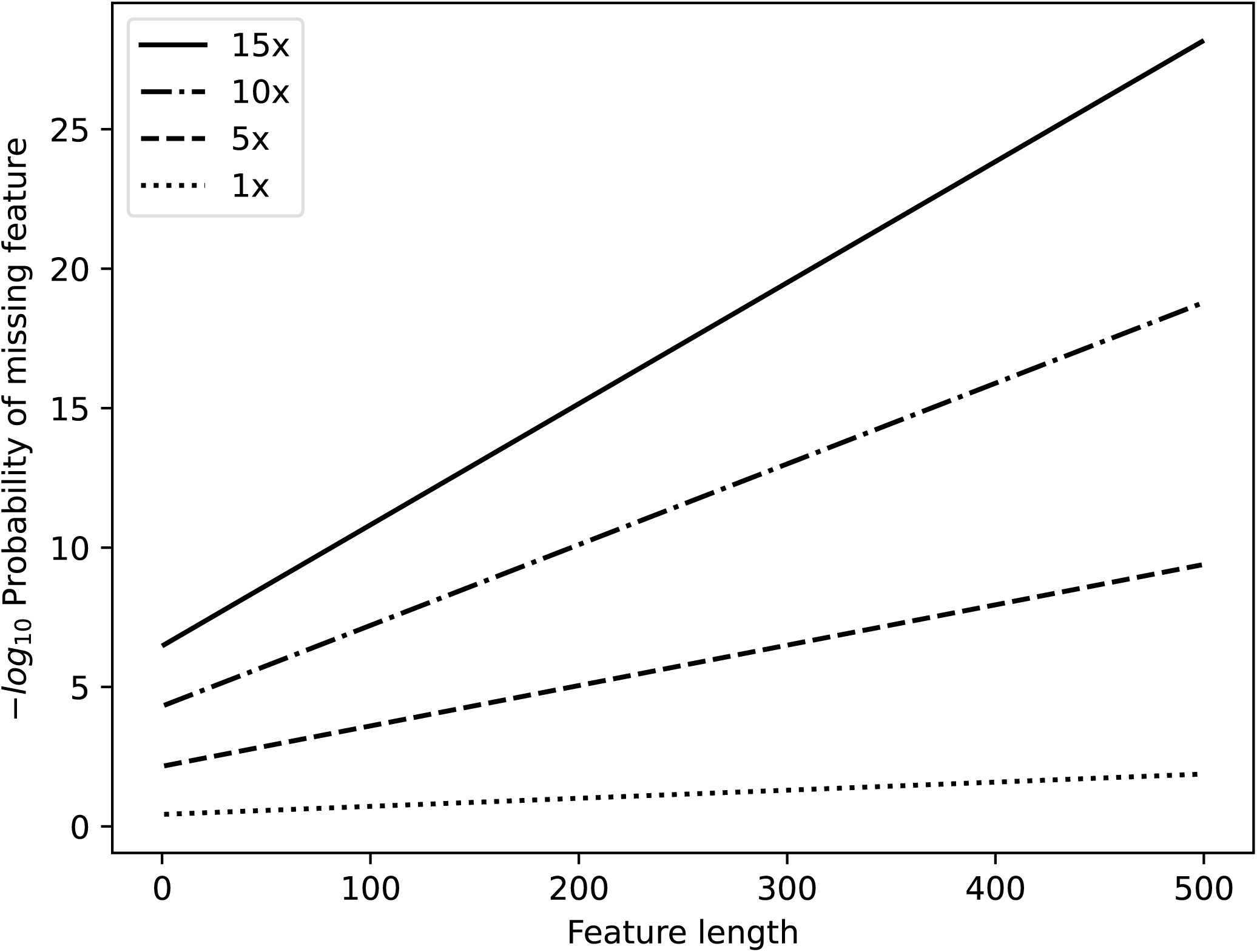
Example plots of the probability of shotgun sequencing missing a genomic feature against feature length for various read-depths (1x, 5x, 10x and 15x). Plotted values were calculated based on 150 bp unpaired reads and a minimum read-feature overlap of 1 bp.

The primary caveat of this result is the first two assumptions, namely that read sampling is unbiased and mapping is perfect. Clearly, deviations from either of these may result in the true value of *Q* going up or down. Nevertheless, when they do hold, even approximately, and read-depth is sufficient, we see that *Q* rapidly vanishes with increasing feature length. Naturally, the specificity required for detecting missing genomic features will vary depending on the research question, however a reasonable rule of thumb seems to be that any average read depth of about 10x and read length of 150 bp will give adequately low (approximately *<<* 10^−5^) values of *Q* for features over about 100 bp.

### A.2 Sequencing quality control

**Figure A.2:**
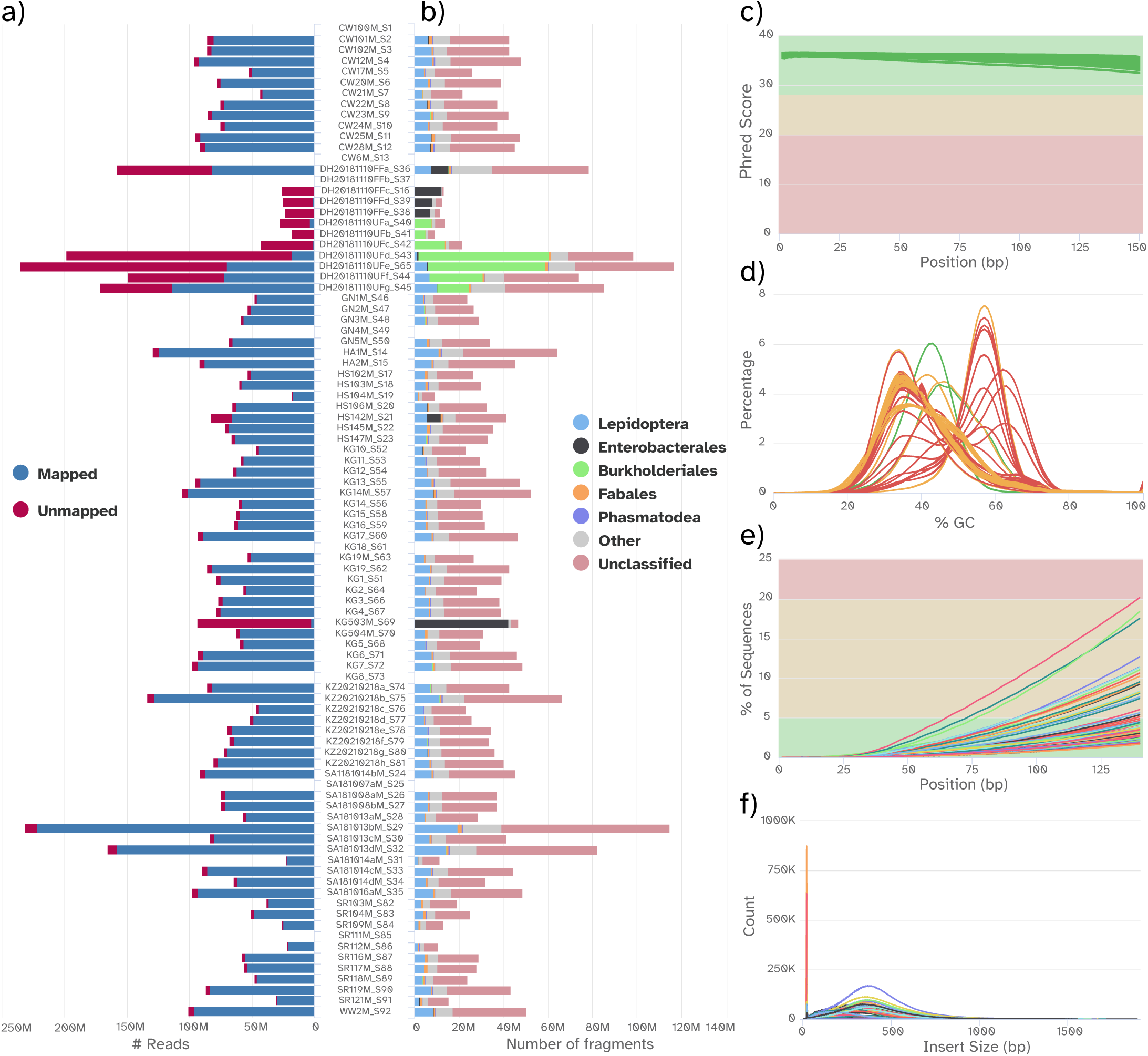
Quality control plots for the whole-genome resequencing experiment. All plots were generated using MultiQC (Ewels et al., 2016). **a**. Total number of reads obtained from each Bogong moth sample, coloured by whether or not the read successfully mapped to the Bogong moth reference genome using BWA-MEM 2 (Vasimuddin et al., 2019). **b**. Number of reads falling into the five most common orders in the dataset, classified by Kraken 2 (Wood et al., 2019) using the NCBI nt database. Non-bogong DNA contamination was present in KG503M and a number of the DH samples (also evident in the mapping rates for those samples shown in **a**). Most reads are unclassified, since the nt database did not include Bogong moth data. **c–e**. Plots of quality-control metrics calculated using FastQC. **c**. Mean quality (Phred score) by read position for each sample. **d**. Distribution of GC content of reads for each sample. Most samples deviated from the expected distribution (red and orange traces). **e**. Sequencing adapter content by read position for each sample. **f**. Distribution of insert sizes of mapped reads, predicted by Picard tools “collectinsertsizemetrics.”

### A.3 Genes under selection

**Table A.1:**
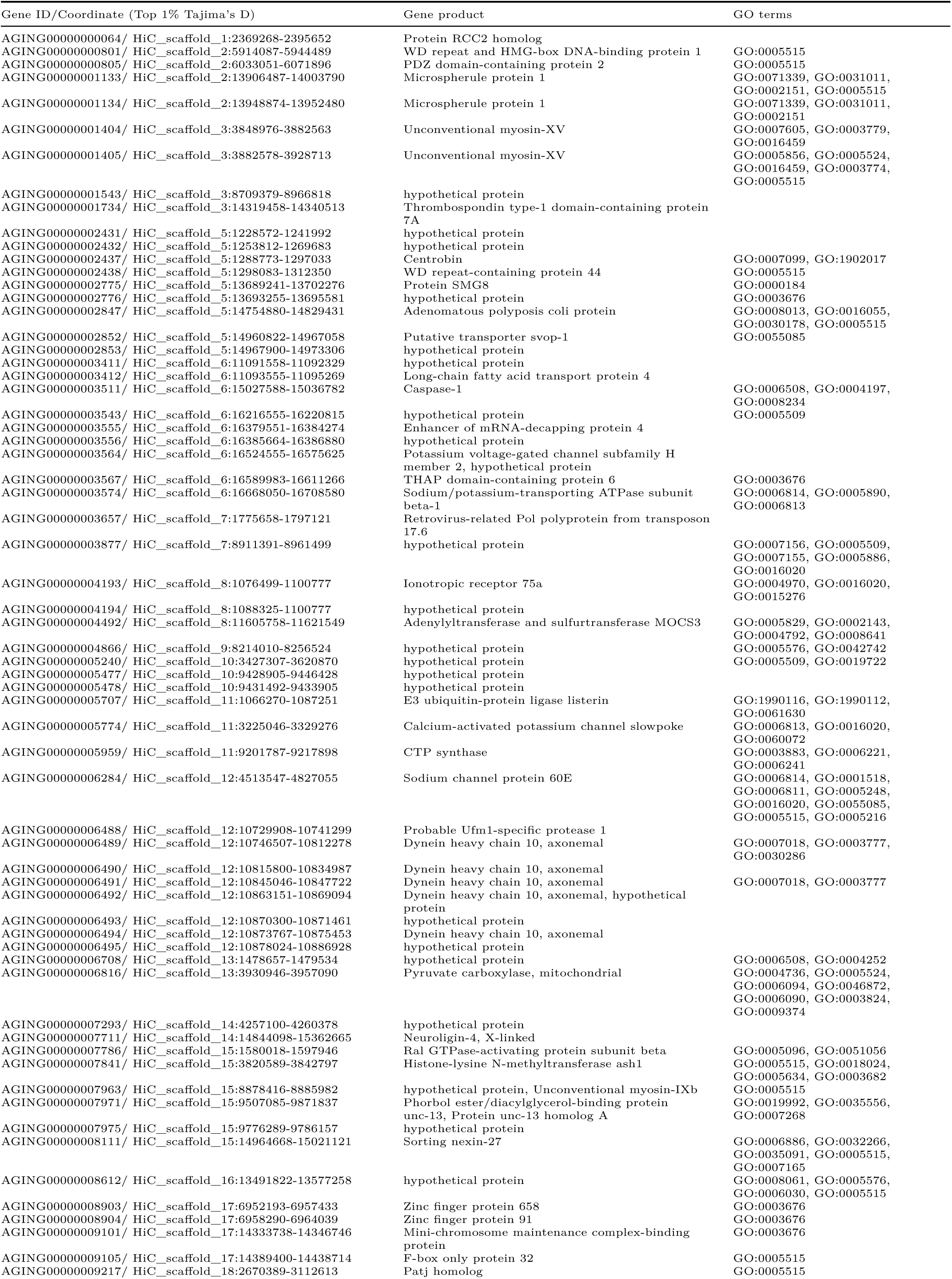
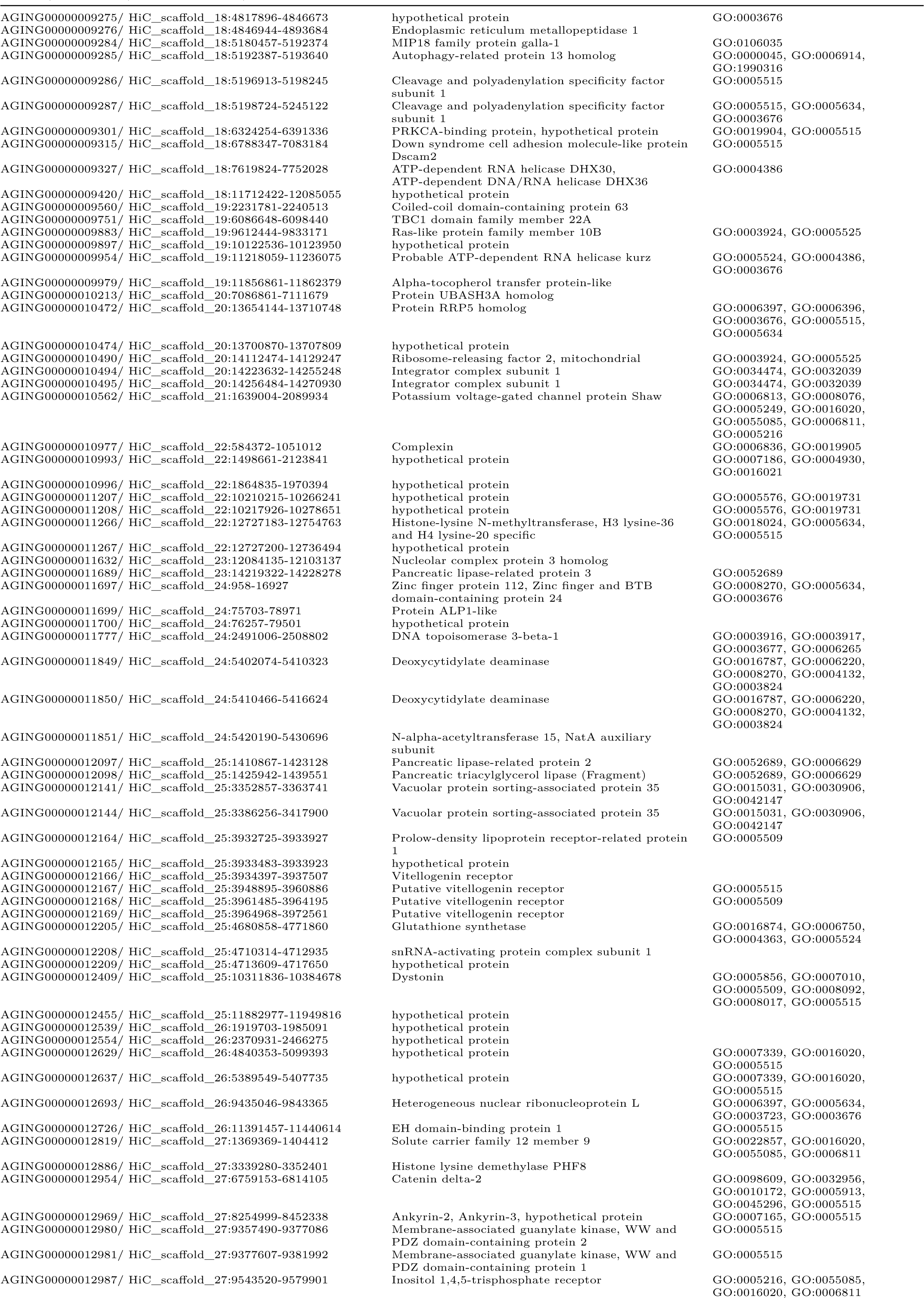
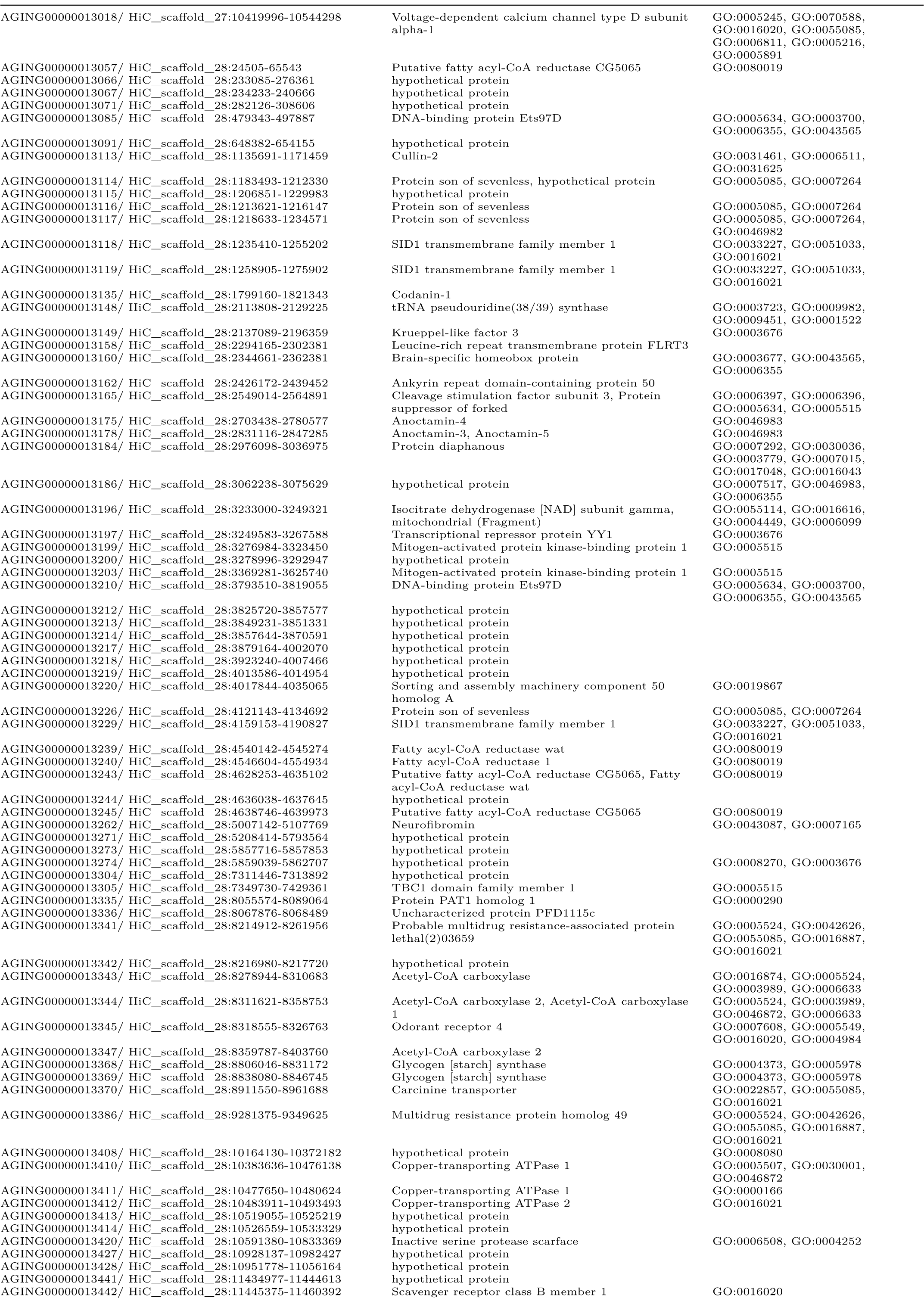
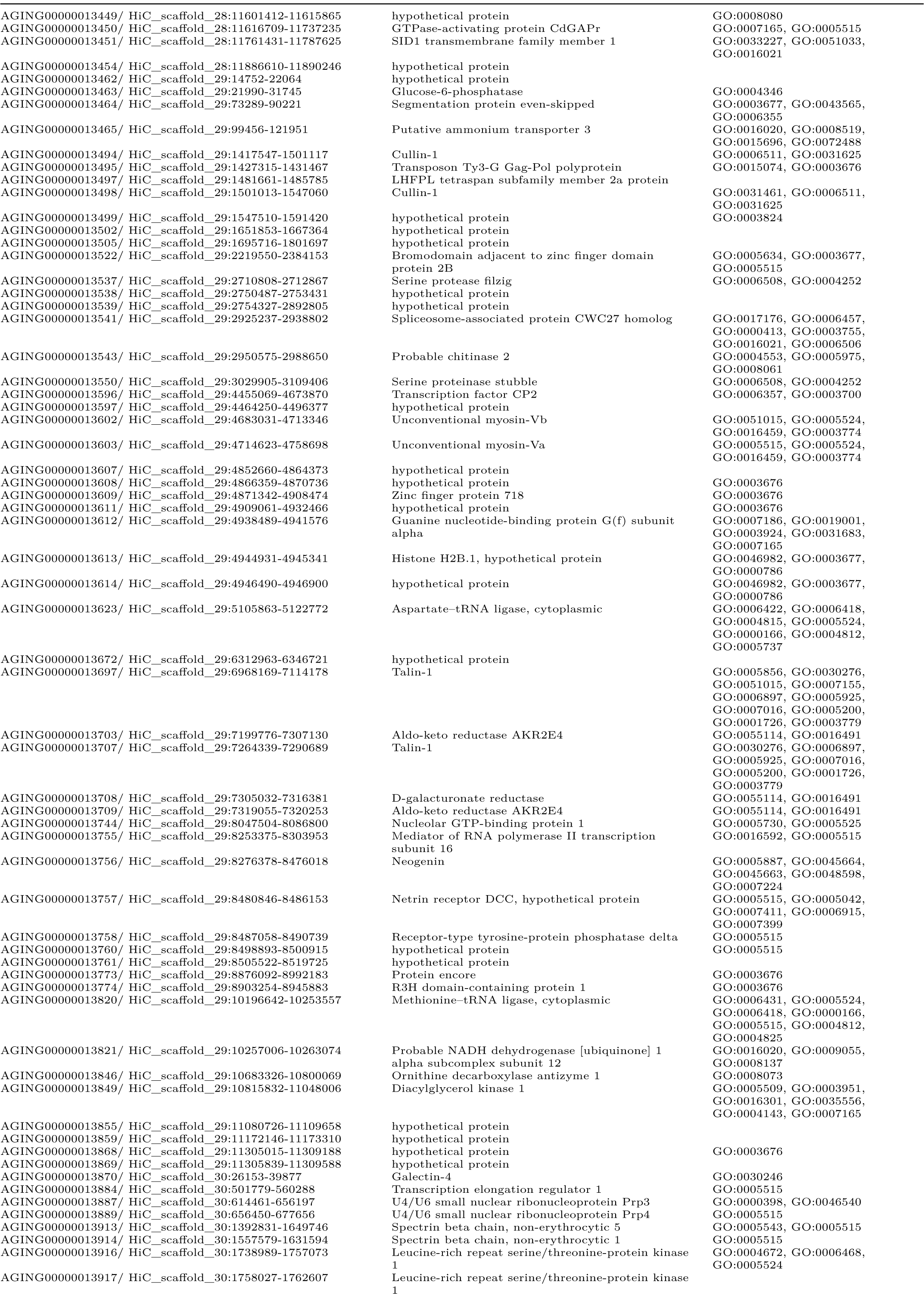
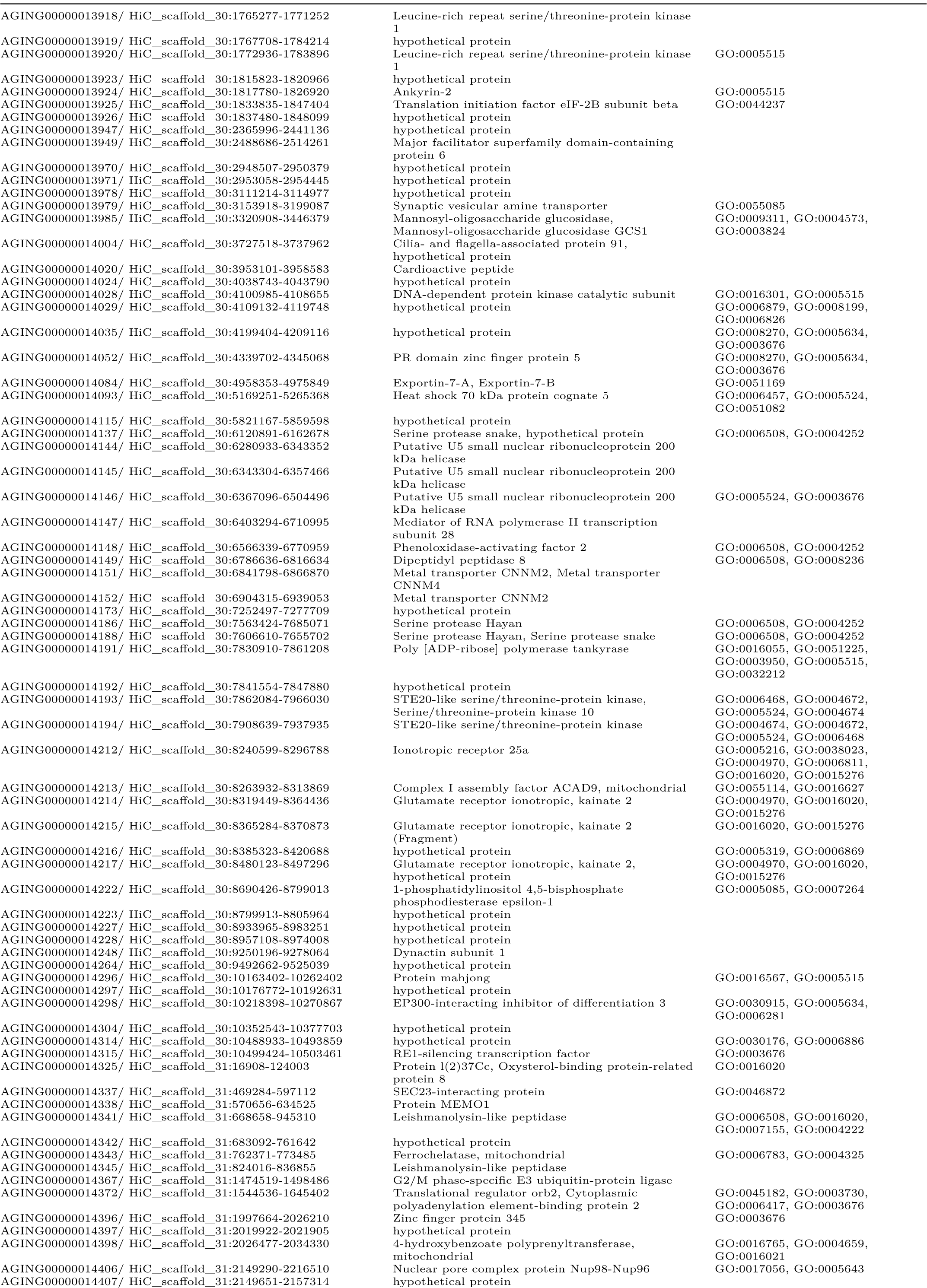
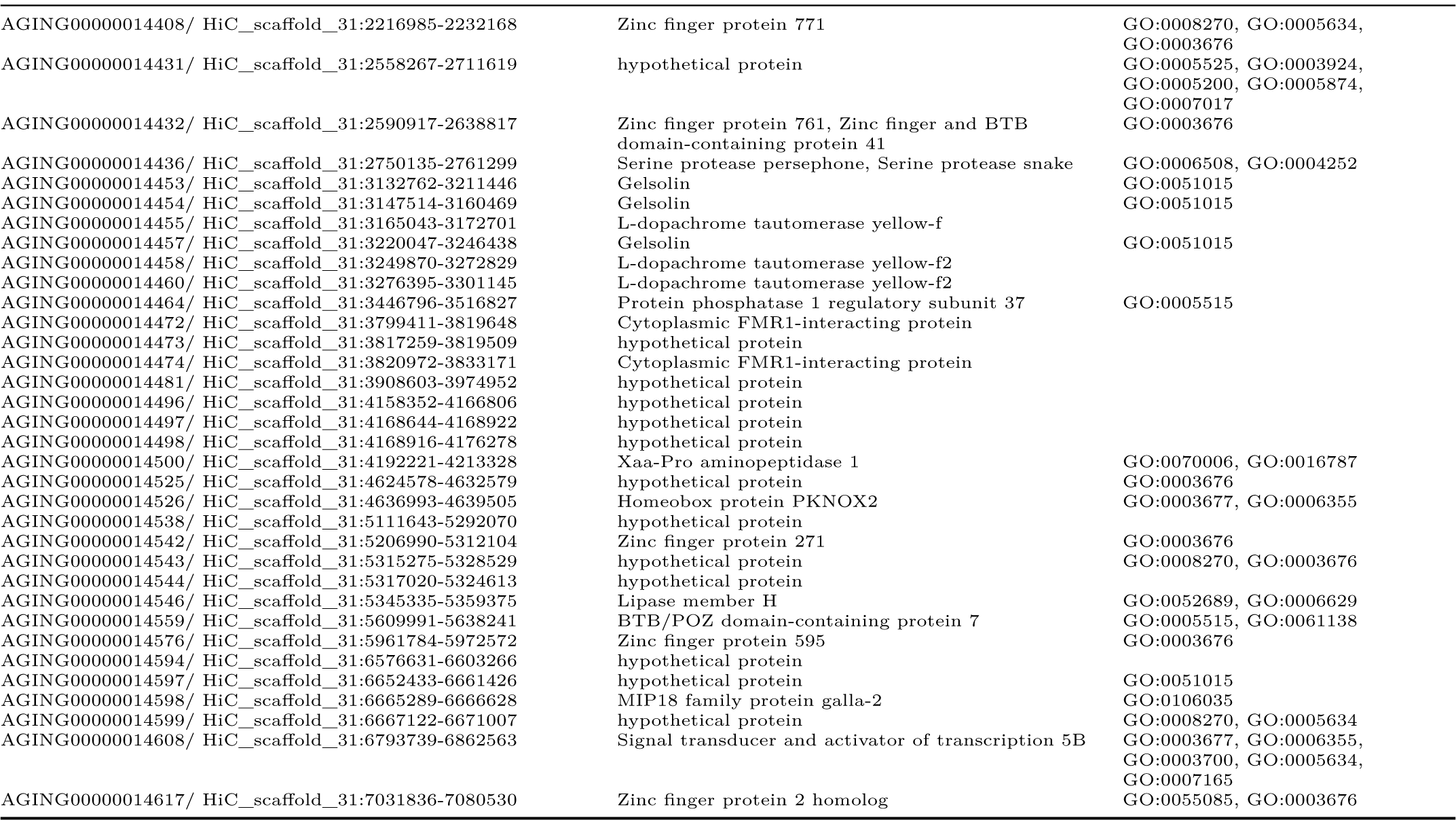
Genes co-located with top 1% of Tajima’s D bins

**Table A.2:**
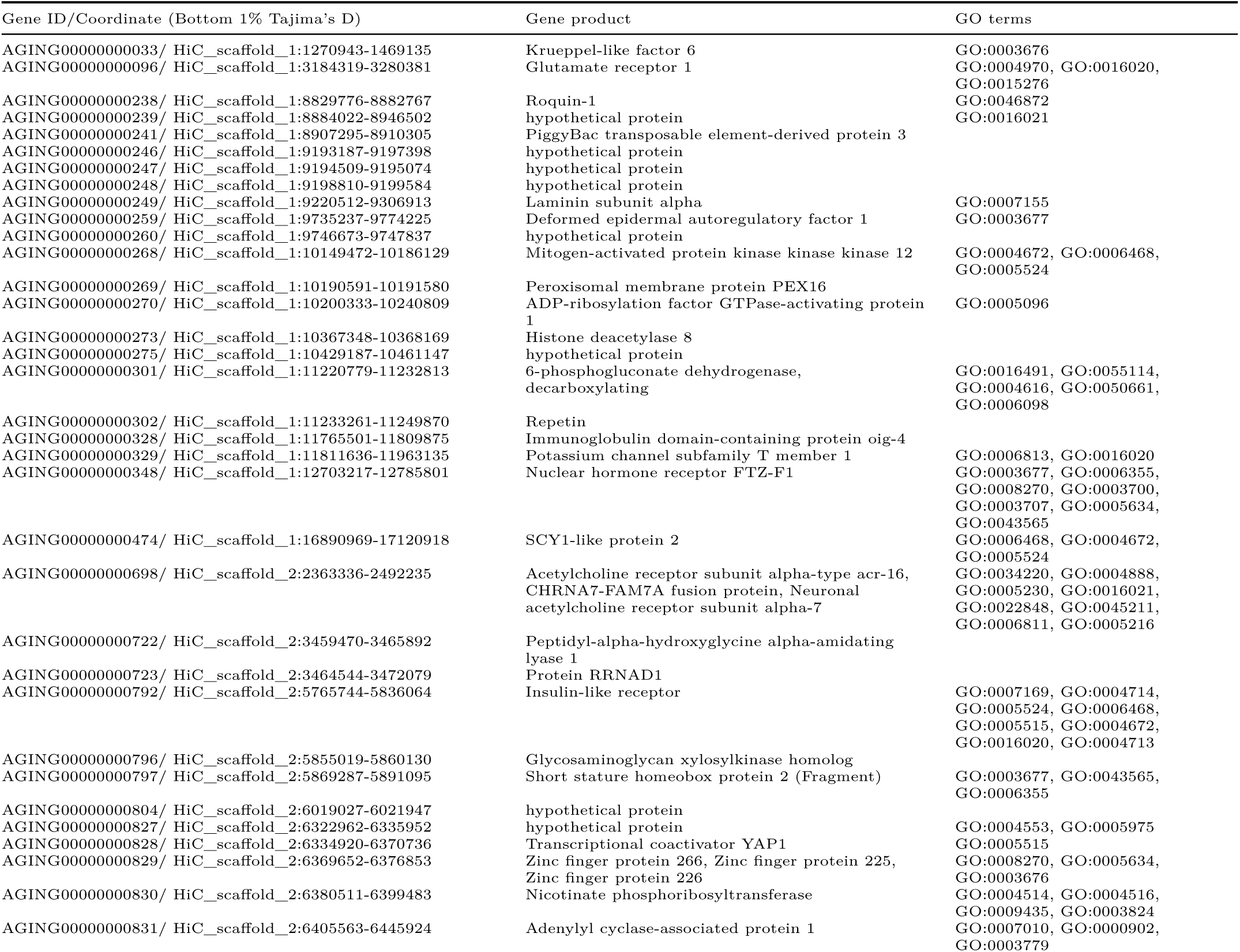
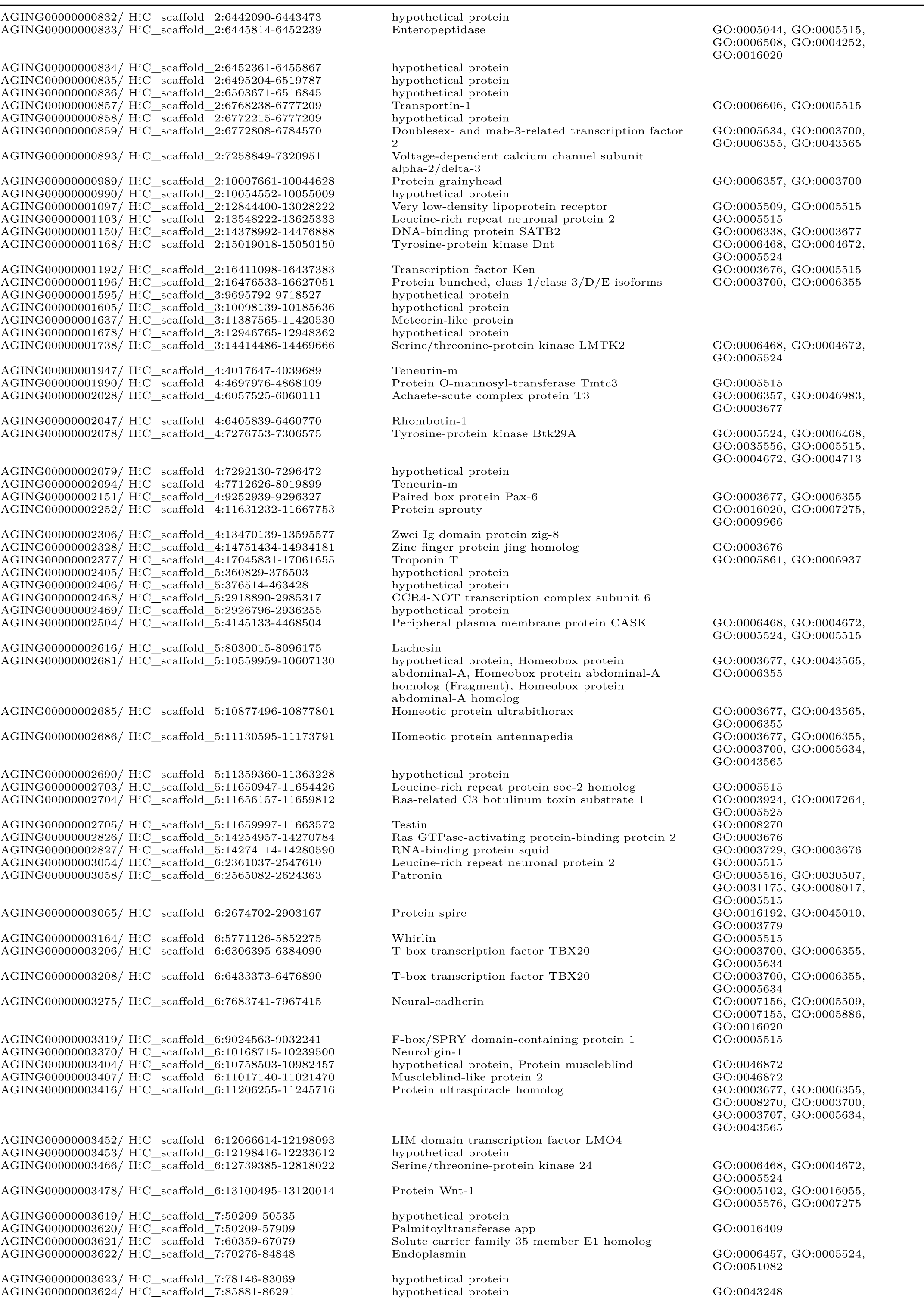
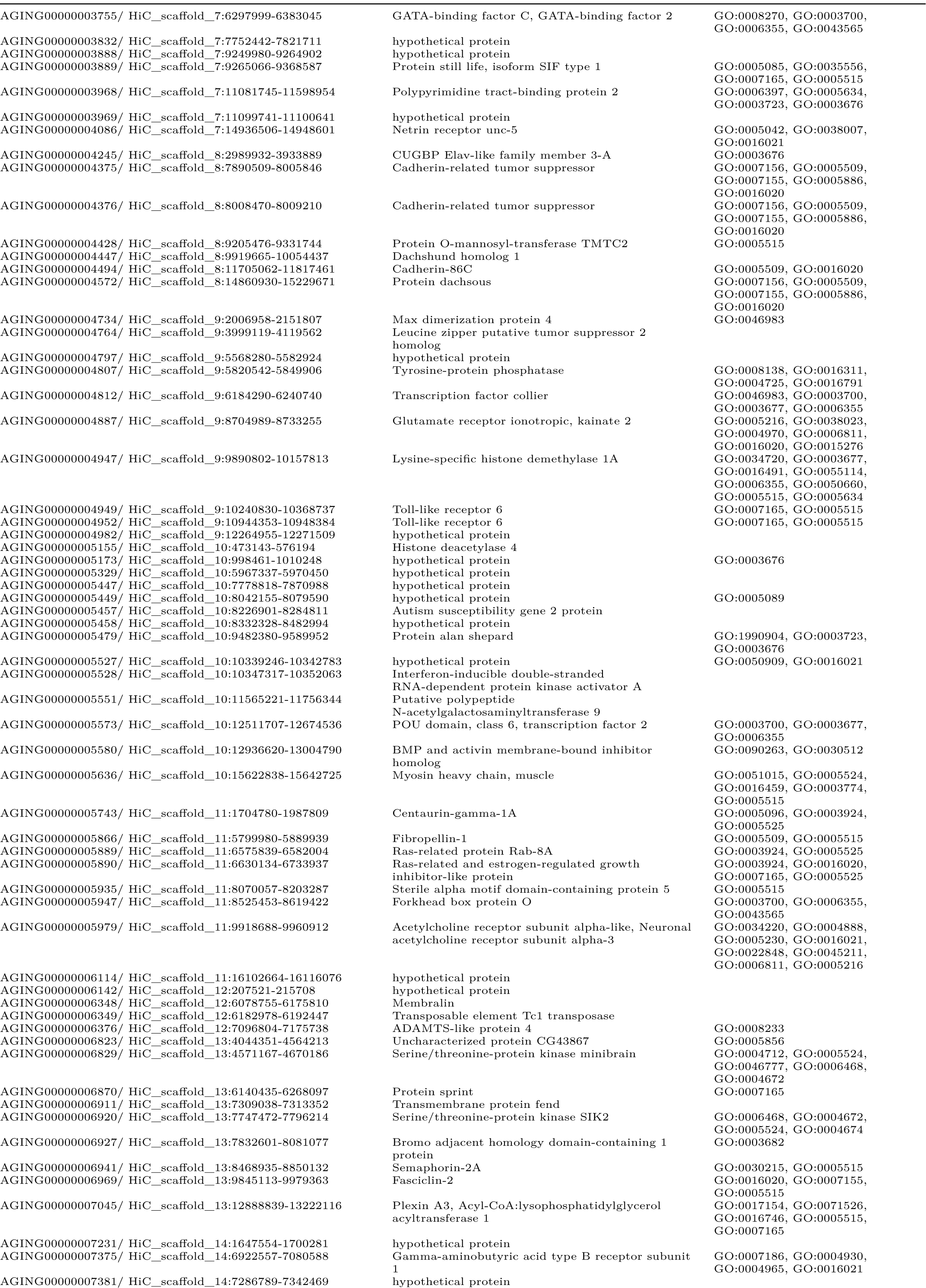
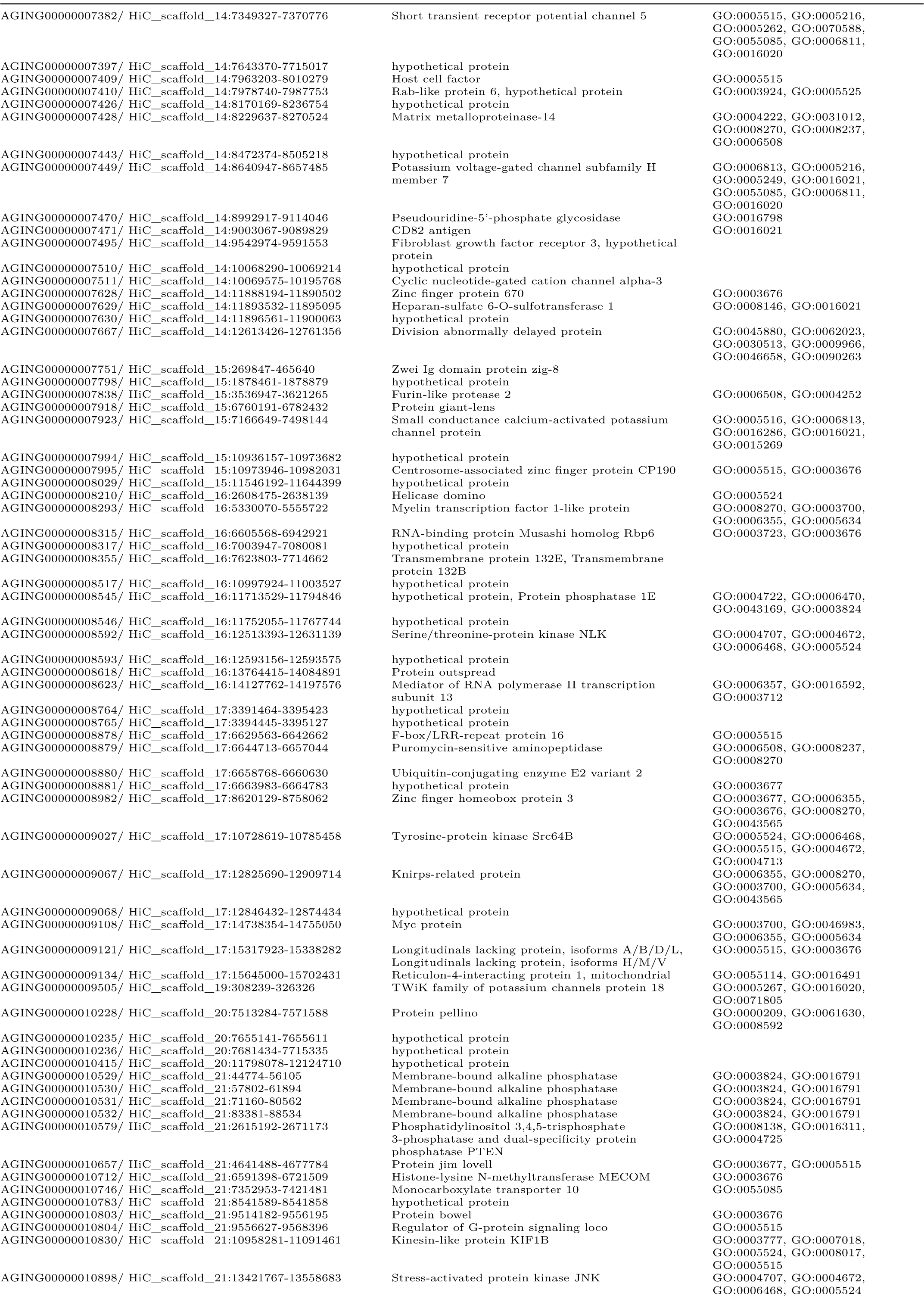
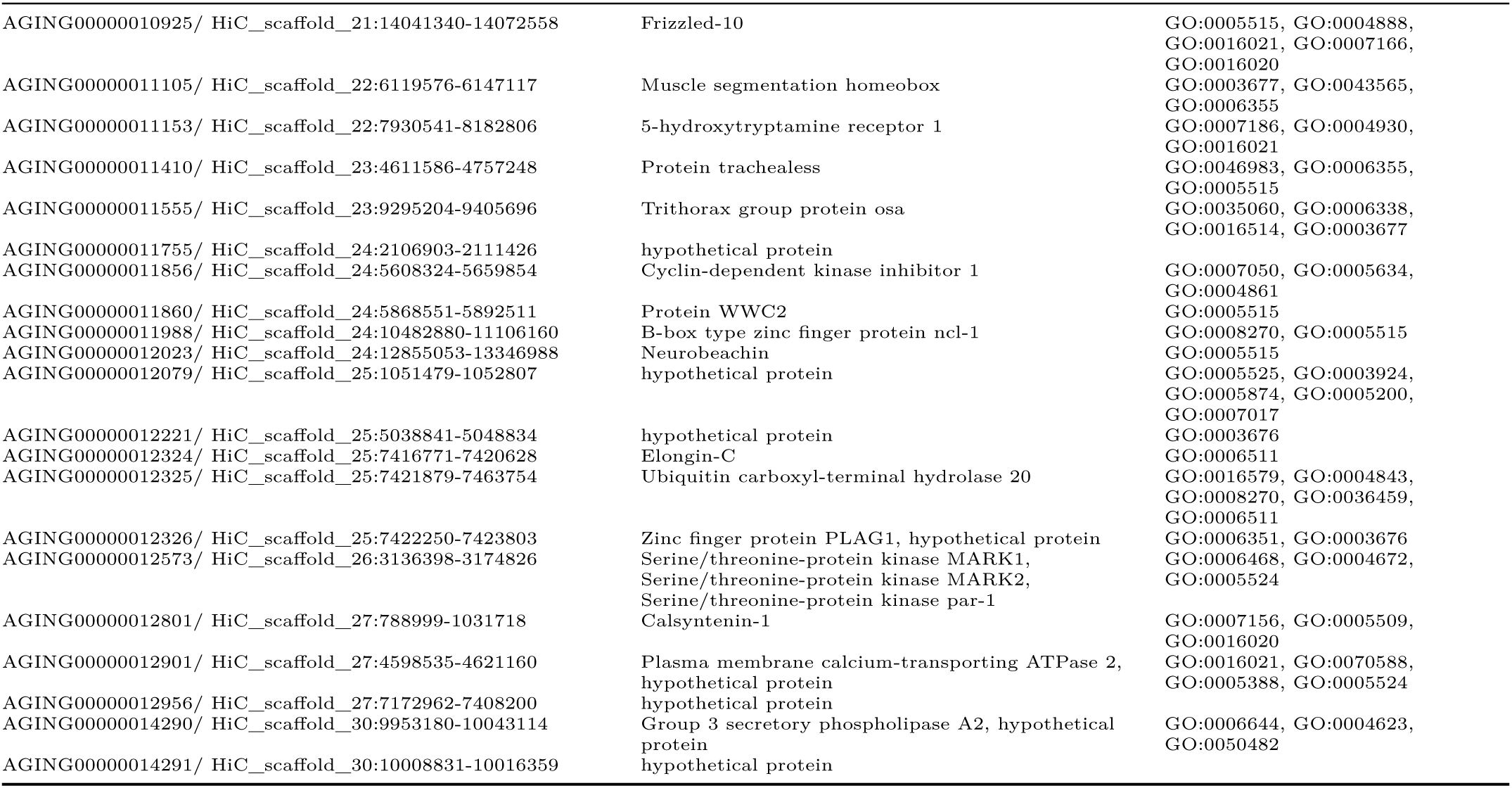
Genes co-located with bottom 1% of Tajima’s D bins

## Footnotes

1 Observations made by Common (1954) at Mt. Gingera led him to conclude that mating does not occur during Bogong moth aestivation. However, in the early hours of the morning on 29^th^ December, 2019, we (JRAW and EJW) observed a pair of Bogong moths copulating in an aestivation site on Mt. Kosciuszko. Whether this is normal or not—merely representing an exception to the usual post-return-migration mating of the moths—remains unclear.

## References

Adden, A., 2020. There and back again: The neural basis of migration in the Bogong moth (PhD thesis). Lund University, Faculty of Science.

Adden, A., Wibrand, S., Pfeiffer, K., Warrant, E., Heinze, S., 2020. The brain of a nocturnal migratory insect, the Australian Bogong moth. Journal of Comparative Neurology 528, 1942–1963. https://doi.org/10.1002/cne.24866

Boisduval, J.A., 1832. Faune entomologique de l’Océan Pacifique: Avec l’illustration des insectes nouveaux recueillis pendant le voyage. J. Tastu.

Bolger, A.M., Lohse, M., Usadel, B., 2014. Trimmomatic: A flexible trimmer for Illumina sequence data. Bioinformatics 30, 2114–2120.

Brehm, G., 2017. A new LED lamp for the collection of nocturnal lepidoptera and a spectral comparison of light-trapping lamps. Nota lepidopterologica 40, 87.

Chaerkady, R., Letzen, B., Renuse, S., Sahasrabuddhe, N.A., Kumar, P., All, A.H., Thakor, N.V., Delanghe, B., Gearhart, J.D., Pandey, A., others, 2011. Quantitative temporal proteomic analysis of human embryonic stem cell differentiation into oligodendrocyte progenitor cells. Proteomics 11, 4007–4020.

Chapman, J.W., Reynolds, D.R., Wilson, K., 2015. Long-range seasonal migration in insects: Mechanisms, evolutionary drivers and ecological consequences. Ecology letters 18, 287–302.

Chen, Q., Yan, W., Duan, E., 2016. Epigenetic inheritance of acquired traits through sperm RNAs and sperm RNA modifications. Nature Reviews Genetics 17, 733–743.

Common, I., 1954. A study of the ecology of the adult bogong moth, Agrotis infusa (Boisd)(Lepidoptera: Noctuidae), with special reference to its behaviour during migration and aestivation. Australian Journal of Zoology 2, 223–263.

Common, I., 1952. Migration and gregarious aestivation in the bogong moth, Agrotis infusa. Nature 170, 981–982.

Common, I.F.B., 1990. Moths of australia. E. J. Brill.

Czech, L., Exposito-Alonso, M., 2021. Grenepipe: A flexible, scalable, and reproducible pipeline to automate variant and frequency calling from sequence reads. arXiv preprint arXiv:2103.15167.

Dingle, H., 2014. Migration: The biology of life on the move. Oxford University Press, USA.

Dreyer, D., Frost, B., Mouritsen, H., Günther, A., Green, K., Whitehouse, M., Johnsen, S., Heinze, S., Warrant, E., 2018. The Earth’s magnetic field and visual landmarks steer migratory flight behavior in the nocturnal Australian Bogong moth. Current Biology 28, 2160–2166. https://doi.org/10.1016/j.cub.2018.05.030

Dreyer, D., Frost, B., Mouritsen, H., Lefèvre, A., Menz, M., Warrant, E., 2021. A guide for using flight simulators to study the sensory basis of long-distance migration in insects. Frontiers in Behavioral Neuroscience.

Ewels, P., Magnusson, M., Lundin, S., Käller, M., 2016. MultiQC: Summarize analysis results for multiple tools and samples in a single report. Bioinformatics 32, 3047–3048.

Farrow, R.A., McDonald, G., 1987. Migration strategies and outbreaks of noctuid pests in australia. International Journal of Tropical Insect Science 8, 531–542.

Friston, K., Tononi, G., Reeke Jr, G., Sporns, O., Edelman, G.M., 1994. Value-dependent selection in the brain: Simulation in a synthetic neural model. Neuroscience 59, 229–243.

Gao, B., Hedlund, J., Reynolds, D.R., Zhai, B., Hu, G., Chapman, J.W., 2020. The ‘migratory connectivity’ concept, and its applicability to insect migrants. Movement Ecology 8, 1–13.

Green, K., 2011. The transport of nutrients and energy into the Australian Snowy Mountains by migrating Bogong moths Agrotis infusa. Austral Ecology 36, 25–34. https://doi.org/10.1111/j.1442-9993.2010.02109.x

Green, K., 2010. The aestivation sites of bogong moths, Agrotis infusa (Boisduval)(Lepidoptera: Noctuidae), in the Snowy Mountains and the projected effects of climate change. Australian Entomologist, The 37, 93–104.

Green, K., Caley, P., Baker, M., Dreyer, D., Wallace, J., Warrant, E., 2021. Australian Bogong moths Agrotis infusa (Lepidoptera: Noctuidae), 1951–2020: Decline and crash. Austral Entomology. https://doi.org/10.1111/aen.12517

Hu, H., Brittain, G.C., Chang, J.-H., Puebla-Osorio, N., Jin, J., Zal, A., Xiao, Y., Cheng, X., Chang, M., Fu, Y.-X., others, 2013. OTUD7B controls non-canonical NF-κB activation through deubiquitination of TRAF3. Nature 494, 371–374.

Hu, H., Wang, H., Xiao, Y., Jin, J., Chang, J.-H., Zou, Q., Xie, X., Cheng, X., Sun, S.-C., 2016. Otud7b facilitates T cell activation and inflammatory responses by regulating Zap70 ubiquitination. Journal of experimental medicine 213, 399–414.

Jackson, T., Wang, H., Nugent, M.J., Griffin, C.T., Burnell, A.M., Dowds, B.C., 1995. Isolation of insect pathogenic bacteria, Providencia rettgeri, from Heterorhabditis spp. Journal of applied bacteriology 78, 237–244.

Keaney, B., others, 2016. Bogong moth aestivation sites as an archive for understanding the Floral, Faunal and Indigenous history of the Northern Australian Alps (PhD thesis). The Australian National University.

Lei, S., He, Z., Chen, T., Guo, X., Zeng, Z., Shen, Y., Jiang, J., 2019. Long noncoding RNA 00976 promotes pancreatic cancer progression through OTUD7B by sponging miR-137 involving EGFR/MAPK pathway. Journal of Experimental & Clinical Cancer Research 38, 1–15.

Li, H., Handsaker, B., Wysoker, A., Fennell, T., Ruan, J., Homer, N., Marth, G., Abecasis, G., Durbin, R., 2009. The sequence alignment/map format and SAMtools. Bioinformatics 25, 2078–2079.

Lundberg, M., Liedvogel, M., Larson, K., Sigeman, H., Grahn, M., Wright, A., Åkesson, S., Bensch, S., 2017. Genetic differences between willow warbler migratory phenotypes are few and cluster in large haplotype blocks. Evolution Letters 1, 155–168.

Martin, M., 2011. Cutadapt removes adapter sequences from high-throughput sequencing reads. EMBnet. journal 17, 10–12.

McKenna, A., Hanna, M., Banks, E., Sivachenko, A., Cibulskis, K., Kernytsky, A., Garimella, K., Altshuler, D., Gabriel, S., Daly, M., others, 2010. The genome analysis toolkit: A MapReduce framework for analyzing next-generation DNA sequencing data. Genome research 20, 1297–1303.

Mölder, F., Jablonski, K.P., Letcher, B., Hall, M.B., Tomkins-Tinch, C.H., Sochat, V., Forster, J., Lee, S., Twardziok, S.O., Kanitz, A., others, 2021. Sustainable data analysis with Snakemake. F1000Research 10.

Murray, D., Zalucki, M., 1990a. Effect of soil moisture and simulated rainfall on pupal survival and moth emergence of Helicoverpa punctigera (Wallengren) and H. Armigera (Hübner)(Lepidoptera: Noctuidae). Australian journal of entomology 29, 193–197.

Murray, D., Zalucki, M., 1990b. Survival of Helicoverpa punctigera (Wallengren) and H. Armigera (Hübner)(Lepidoptera: Noctuidae) pupae submerged in water. Australian journal of entomology 29, 191–192.

Müssig, C., Schröder, F., Usadel, B., Lisso, J., 2010. Structure and putative function of NFX1-like proteins in plants. Plant Biology 12, 381–394.

O’Donnell, K.J., Meaney, M.J., 2020. Epigenetics, development, and psychopathology. Annual Review of Clinical Psychology 16, 327–350.

Pina-Martins, F., Silva, D.N., Fino, J., Paulo, O.S., 2017. Structure_threader: An improved method for automation and parallelization of programs STRUCTURE, fastStructure and MavericK on multicore CPU systems. Molecular ecology resources 17, e268–e274.

Raj, A., Stephens, M., Pritchard, J.K., 2014. fastSTRUCTURE: Variational inference of population structure in large SNP data sets. Genetics 197, 573–589.

Rittschof, C.C., Bukhari, S.A., Sloofman, L.G., Troy, J.M., Caetano-Anollés, D., Cash-Ahmed, A., Kent, M., Lu, X., Sanogo, Y.O., Weisner, P.A., others, 2014. Neuromolecular responses to social challenge: Common mechanisms across mouse, stickleback fish, and honey bee. Proceedings of the national Academy of Sciences 111, 17929–17934.

Sims, S.R., 2008. Influence of soil type and rainfall on pupal survival and adult emergence of the fall armyworm (Lepidoptera: Noctuidae) in southern Florida. Journal of Entomological Science 43, 373–380.

Sporns, O., Almássy, N., Edelman, G.M., 2000. Plasticity in value systems and its role in adaptive behavior. Adaptive Behavior 8, 129–148.

Stephenson, B., David, B., Fresløv, J., Arnold, L.J., Delannoy, J.-J., Petchey, F., Urwin, C., Wong, V.N., Fullagar, R., Green, H., others, 2020. 2000 year-old Bogong moth (Agrotis infusa) Aboriginal food remains, Australia. Scientific reports 10, 1–10.

Tajima, F., 1989. Statistical method for testing the neutral mutation hypothesis by DNA polymorphism. Genetics 123, 585–595.

Talla, V., Pierce, A.A., Adams, K.L., Man, T.J. de, Nallu, S., Villablanca, F.X., Kronforst, M.R., Roode, J.C. de, 2020. Genomic evidence for gene flow between monarchs with divergent migratory phenotypes and flight performance. Molecular ecology 29, 2567–2582.

Tenger-Trolander, A., Lu, W., Noyes, M., Kronforst, M.R., 2019. Contemporary loss of migration in monarch butterflies. Proceedings of the National Academy of Sciences 116, 14671–14676.

Vasimuddin, M., Misra, S., Li, H., Aluru, S., 2019. Efficient architecture-aware acceleration of BWA-MEM for multicore systems, in: 2019 IEEE International Parallel and Distributed Processing Symposium (IPDPS). IEEE, pp. 314–324.

Villanueva, P., Nudel, R., Hoischen, A., Fernández, M.A., Simpson, N.H., Gilissen, C., Reader, R.H., Jara, L., Echeverry, M.M., Francks, C., others, 2015. Exome sequencing in an admixed isolated population indicates NFXL1 variants confer a risk for specific language impairment. PLoS genetics 11, e1004925.

Wang, B., Jie, Z., Joo, D., Ordureau, A., Liu, P., Gan, W., Guo, J., Zhang, J., North, B.J., Dai, X., others, 2017. TRAF2 and OTUD7B govern a ubiquitin-dependent switch that regulates mTORC2 signalling. Nature 545, 365–369.

Warrant, E., Frost, B., Green, K., Mouritsen, H., Dreyer, D., Adden, A., Brauburger, K., Heinze, S., 2016. The Australian Bogong moth Agrotis infusa: A long-distance nocturnal navigator. Frontiers in behavioral neuroscience 10. https://doi.org/10.3389/fnbeh.2016.00077

Warrant, E.J., Whitehouse, M.E.A., Green, K., Wallace, J.R.A., Caley, P., Tomlinson, S., Umbers, K., 2021. Agrotis infusa. The IUCN Red List of Threatened Species 2021. https://doi.org/10.2305/IUCN.UK.2021-3.RLTS.T190513532A196183274.en

Wedd, L., Kucharski, R., Maleszka, R., 2016. Differentially methylated obligatory epialleles modulate context-dependent LAM gene expression in the honeybee Apis mellifera. Epigenetics 11, 1–10.

Welch, H., 1963. Amphimermis bogongae sp. Nov. And Hexamermis cavicola sp. Nov. From the Australian bogong moth, Agrotis infusa (Boisd.), With a review of the genus Amphimermis Kaburaki & Imamura, 1932 (Nematoda: Mermithidae). Parasitology 53, 55–62.

Wood, D.E., Lu, J., Langmead, B., 2019. Improved metagenomic analysis with Kraken 2. Genome biology 20, 1–13.

Xu, J., Jarocha, L.E., Zollitsch, T., Konowalczyk, M., Henbest, K.B., Richert, S., Golesworthy, M.J., Schmidt, J., Déjean, V., Sowood, D.J., others, 2021. Magnetic sensitivity of cryptochrome 4 from a migratory songbird. Nature 594, 535–540.

You, M., Ke, F., You, S., Wu, Z., Liu, Q., He, W., Baxter, S.W., Yuchi, Z., Vasseur, L., Gurr, G.M., others, 2020. Variation among 532 genomes unveils the origin and evolutionary history of a global insect herbivore. Nature communications 11, 1–8.

Zhou, X., Stephens, M., 2012. Genome-wide efficient mixed-model analysis for association studies. Nature genetics 44, 821–824.

